# Heat tolerance and canopy temperatures of *Larix sibirica* under highly continental climate in Mongolia’s boreal forest

**DOI:** 10.64898/2026.06.30.735460

**Authors:** Choimaa Dulamsuren, Jayamaran Temuulen Abbas, Germar Csapek, Erdenechuluun Naranbayar, Tumendelger Uitumen, Dugersuren Amarjargal, Gurbazar Byamba-Yondon, Davaadorj Saindovdon, Tungalag Munkhzul, Ganbaatar Batsaikhan, Markus Hauck

## Abstract

Compared to drought stress, direct heat damage has been considered secondary as a cause of climate change-induced tree mortality and productivity declines in forests. However, evidence from temperate, subtropical and tropical forests is accumulating that direct heat damage in photosystem II (PS II) is also a realistic scenario under climate change. We analyzed PS II heat tolerance in *Larix sibirica*, which represents a dominant boreal tree species in Siberia and northern Central Asia in cold environments with subzero or near-zero mean annual temperatures, but nevertheless warm summers. Thermal imaging was applied to relate heat thresholds found in the laboratory to canopy temperatures. Measurements were repeated in three months during the short growing season to test for the occurrence of heat acclimation.

After 4 h of heat exposure, the temperature of the most rapid decline of the maximum fluorescence yield (*F*_v_/*F*_m_) due to heat (*T*_IP_), which is thought to indicate irreversible damage, was 41.2±0.0°C. The critical temperature (*T*_crit_) indicating initial reversible heat stress in PS II was 39.8±0.1°C. *L. sibirica* had large thermal safety margins, when *T*_IP_ and *T*_crit_ were compared to the canopy temperatures measured during the study period, but not compared to record heat maxima since 2000. The trees acclimated to heat over summer by increasing *T*_IP_ and thus increasing the tolerance to severe heat stress. However, tolerance to moderate heat (indicated by *T*_crit_) was simultaneously weakened, suggesting the reallocation of resources during acclimation. In most summers, *L. sibirica* forests are not threatened by direct heat damage, but heat extremes as previously recorded from our study region in Mongolia’s boreal forest could increase mortality due to PS II damage. Heat can thus be seen as one out of several stressors that reduces the vitality and growth of southern boreal forests with *L. sibirica* under climate change.

## 1. Introduction

The increased occurrence and intensification of hot droughts due to climate change has globally led to increased tree mortality (Allen et al. 2010; Hammond et al. 2022) and partly to reductions in annual stem increment (Anderson-Teixeira et al. 2022; Mirabel et al. 2023). Often reduced increment and increased growth variability precedes increased mortality (Cailleret et al. 2019; Dulamsuren et al. 2022; Keen et al. 2022). Research on the ecophysiology of climate change-induced growth declines and increases in tree mortality had a strong focus on tree water relations, including studies of hydraulic failure, dehydration of sensitive tissues, and carbon starvation as the result of stomatal closure that increased to avoid critical water losses (Mantova et al. 2021, 2023). Promotion of pathogens (Caldeira 2019), herbivores (Anderegg et al. 2015), and wildfire (Cansler et al. 2024) have also been extensively studied.

Direct effects of heat stress on trees have been less well studied, because drought stress was considered more relevant to trigger tree mortality and reduced growth under climate change. Though already Rennenberg et al. (2006) noted that trees might suffer from heat damage in the photosystem II (PS II) during hot droughts, heat was mostly seen as a threat for trees due to its indirect influence on tree water relations by increasing the atmospheric vapor pressure deficit (VPD) and reducing soil moisture (Allen et al. 2010; Breshears et al. 2013; Williams et al. 2013). Nevertheless, evidence is accumulating which suggests that thresholds for heat damage in PS II can be crossed during heat waves in the field in both temperate and tropical latitudes (Tiwari et al. 2021; Hauck et al. 2025). Studies on the heat tolerance of tree species from boreal forests, where physiological thresholds for heat damage are related to canopy temperatures in the field are lacking, so far. In temperate tree species, high tolerance to heat was found to be correlated with high drought tolerance in many species, but there are notable exceptions of drought-tolerant species with moderate or low heat tolerance (Hauck et al. 2025; Kunert et al. 2026a).

The principle mechanisms of heat stress in vascular plants are generally well-studied (Allakhverdiev et al. 2008; Mohanty et al. 2012). The photosystem II (PS II) plays a special role in heat tolerance, because it responds particularly sensitively to heat and because damage can be precisely detected with chlorophyll fluorescence analysis. PS II dysfunction is caused by the dissociation of the oxygen-evolving complex, of the chlorophyll *a*-containing core complex into two protein monomers, and of the light-harvesting complexes from the core complex (Yamamoto et al. 2008; Lípová et al. 2010). Increased fluidity of thylakoids as of other phospholipid membranes and grana destacking in the thylakoid membranes also contribute to PS II dysfunction under heat (Yamamoto 2016). Reactive oxygen species (ROS), which are produced when the photochemical water splitting process at the oxygen-evolving complex is disturbed, are an additional source of heat damage (Yamashita et al. 2008; Pospíšil 2016).

There are repair mechanisms for PS II damage, but unfortunately these repair mechanisms are more sensitive to heat than the PS II itself (Mohanty et al. 2012). This situation, which at first glance seems paradoxical, is attributable to the fact that PS II repair is also necessary as a response to photoinhibition, which is more common than heat damage (Murata et al. 2007). Photosystem I (PS I) is more heat-stable than PS II, which makes PS II to a primary target of heat damage (Berry & Björkman 1980). The dark reaction of photosynthesis is also sensitive to heat due to inactivation of Rubisco activase, but this effect is fully reversible, as the enzyme is not disintegrated (Salvucci & Crafts-Brandner 2004). By the production of heat shock proteins (HSP) and alterations in the membrane lipid composition, plants can acclimate to heat (Posch et al. 2025). Heat acclimation during the growing season has been suggested for temperate broadleaves and conifers (Húdoková et al. 2022; Hauck et al. 2026).

How realistically laboratory experiments on the heat tolerance of trees reflect heat stress in the field depends, among others, on canopy temperatures. Most laboratory experiments examine the effect of heat in detached leaves or even leaf disks (Krause et al., 2010; Kunert et al., 2022; Münchinger et al. 2023; Da Silva & Rossatto 2024). Therefore, such experiments, including the present ones, refer to given leaf temperatures, but do not capture how trees can mitigate heating of their foliage during heat waves by transpirational cooling (Gauthey et al. 2024). Transpiration links tree water relations with heat tolerance and contributes to the temperature gap between the leaf and the atmosphere (Yi et al. 2020). Canopy and leaf morphology can also affect leaf temperatures (Still et al. 2021). While the precise measurement of canopy temperatures was formerly an intricate endeavor (Leuzinger & Körner 2007), it has become much easier since drone-borne thermal imaging has been developed (Still et al. 2019; Richter et al. 2021; van Tiel et al. 2026). Nonetheless, studies that directly relate laboratory experiments on heat tolerance to canopy temperatures in the field are scarce (Endris & Rehm 2025). Mostly leaf temperatures exceed air temperatures (Still et al. 2022).

The relationship between canopy temperature and the physiological heat tolerance of trees is expressed as the thermal safety margin (TSM), where maximum temperatures from the field are related to heat tolerance indices determined during heat exposure experiments in the laboratory (O’Sullivan et al. 2017; Perez & Feeley 2020). There are different definitions for heat tolerance indices that quantify heat damage in plants. We rely most on the temperature (*T*_IP_) and the *F*_v_/*F*_m_ value (*F*_IP_) at the inflection point of the *F*_v_/*F*_m_-vs.-temperature function, which is a sigmoid relationship (Hauck & Dulamsuren 2026). This metric was recently proposed by Hauck & Dulamsuren (2026), because it identifies the point of the most rapid decline in PS II functioning and is thus highly likely to represent a mechanistically relevant temperature of severe PS II damage. It was suggested to use *T*_IP_ and *F*_IP_ to replace the similar absolute (*T*_50_) and relative (*T*_50_’) versions of the temperature of 50% *F*_v_/*F*_m_ reduction, because these metrics have less explanatory power and definitions of *T*_50_ in the past were inconsistent (Hauck & Dulamsuren 2026). *T*_50_ specifies the temperature of 50% *F*_v_/*F*_m_ reduction without considering whether this temperature is in a curve section of rapid *F*_v_/*F*_m_ decline or not (Krause et al. 2010) and is thus not directly supported by a mechanism. *T*_50_’ attempts to avoid this problem by standardizing *F*_v_/*F*_m_ from 0 to 1 (Tiwari et al. 2021), but that way all information on the severity of the *F*_v_/*F*_m_ decline under heat is lost, which is, however, highly relevant for assessing the reversibility of damage (Hauck & Dulamsuren 2026). In addition to *T*_IP_ and *F*_IP_, we use the critical temperature (*T*_crit_), which represents the temperature of initial *F*_v_/*F*_m_ decline (Hüve et al. 2011; Perez & Feeley 2020) and can be assumed to represent membrane damage to an extent that is largely reversible (Kreslavski et al. 2008; Didion-Gency et al. 2025).

In the present study, we analyzed the heat tolerance of *Larix sibirica* as a case example of an ecologically and quantitatively important boreal trees, combining sampling for laboratory experiments and thermal imaging from the same forest stands. We also included the potential for heat acclimation in our study by repeating the measurements during summer from June to August. *L. sibirica* forests from the southern edge of the boreal forest biome in Mongolia were selected as a model system for our study, because most of these forests are naturally drought-limited, face strongly increased temperatures due to climate change and respond with reduced annual stem increment and increased tree mortality in the recent past (Dulamsuren et al. 2010; Liu et al. 2013; Dulamsuren 2026). While ecophysiological and dendrochronological research has suggested increased drought as a key factor for this development, the objective of the present study was to investigate whether heat acts as an additional stressor that contributes to the reduced growth and increased mortality of *L. sibirica*.

Based on existing measurements of other conifers of the Pinaceae from the northern hemisphere (Hauck et al. 2025), we hypothesized (1) that the PS II of *L. sibirica* is relatively heat-stable at temperatures of 35–40°C, but is sensitive to temperatures beyond this threshold. Since Petrik et al. (2023) and Hauck et al. (2025) partly found increased *F*_v_/*F*_m_ under heat in summer than in spring in other Pinaceae (*Abies alba*, *Picea abies*), which could be interpreted as a sign of heat acclimation during the summer, we tested the hypothesis (2) that *L. sibirica* increasingly acclimates its PS II to heat from June to August. Due to their position at the southern edge of the boreal forest in the ecotone to the Central Asian steppe, Mongolia’s *L. sibirica* forests are mostly limited to the moister and cooler north-facing slopes, whereas south-facing slopes are covered with grassland (Klinge et al. 2018). Therefore, we limited our studies to north-facing mountain slopes as the most typical habitat of *L. sibirica* at its southern distribution limit. Because of the generally low temperatures in Mongolia (Dulamsuren & Hauck 2008), we hypothesized (3) that canopy temperature in the *L. sibirica* forests on north-facing mountain slopes are usually below the threshold for severe heat damage (as expressed by TSM) despite of climate warming.

## 2. Materials and methods

### 2.1. Study area

The study was conducted at the southern edge of the Central Asian/southern Siberian boreal forest region in the Khangai Mountains, northern-central Mongolia near the town of Jargalant (48°44’ N, 100°46’ E, 1600 m a.s.l.), Arkhangai district. Forests in the area are strongly dominated by *Larix sibirica* Ledeb. and mostly limited to north-facing mountains slopes, whereas south-facing slopes are too dry for forest and usually covered with grassland. There is no systematic forest management in the region, but forests are sporadically used for selective logging of dead trees.

### 2.2. Climate of the study area

The climate of Mongolia’s boreal forests is highly continental with cold, dry winters caused by the Siberian anticyclone, subzero mean annual temperatures at many places, and a short summer with variable precipitation that are received when the Siberian anticyclone weakens and gives way to the inflow of moist airmasses from the westerlies. Long-term climate data for our study region was obtained as modeled data (1901–2023) from the CRU TS 4.08 dataset (Climate Research Unit of the University of East Anglia, Norwich and Met Office, Exeter, UK; https://climexp.knmi.nl) at a resolution of 0.5° × 0.5° (Harris et al. 2020) for the grid field of 48.5 – 49.0 °N and 100.5 – 101.0 °E. As we conducted our field work in summer 2025, this dataset was extended with Copernicus ERA5 reanalysis data (https://cds.climate.copernicus.eu).

Mean annual air temperature (MAT) for 1901–2023 was –3.1°C with lower and upper 5% confidence intervals of –3.2°C and –2.9°C. Mean July and January temperatures amounted to 13.7°C and –21.7°C. Temperatures have increased since 1901, with an intensification after the 1970s, as is displayed for MAT in Fig. 1a and for July temperatures in Fig. 1b. Based on linear regression, MAT has increased by 0.22 K decade^−1^ (or a total of 2.7 K) from 1901–2023 and by 0.68 K decade^−1^ (or 2.3 K) from 1970–2023. The increase in temperature concerned all seasons (Fig. 1c). The highest temperatures during the hottest month July in the ERA5 dataset (2000–2025) occurred on 19 JUL 2002 with a maximum of 33.0°C and on 20 JUL 2002 with a maximum of 32.6 °C. Temperatures ≥30°C lasted for 7 h on the first day of this heat wave and for 9 h on the second day. At the nearest instrumental weather station in Erdenemandal (48°31’ N, 101°22’ E, 1509 m a.s.l.; ca. 50 km SE of our study area), where also *L. sibirica* forests occur, a daily maximum temperature of 35.6°C was recorded in 2002 (https://climexp.knmi.nl).

**Fig. 1.**
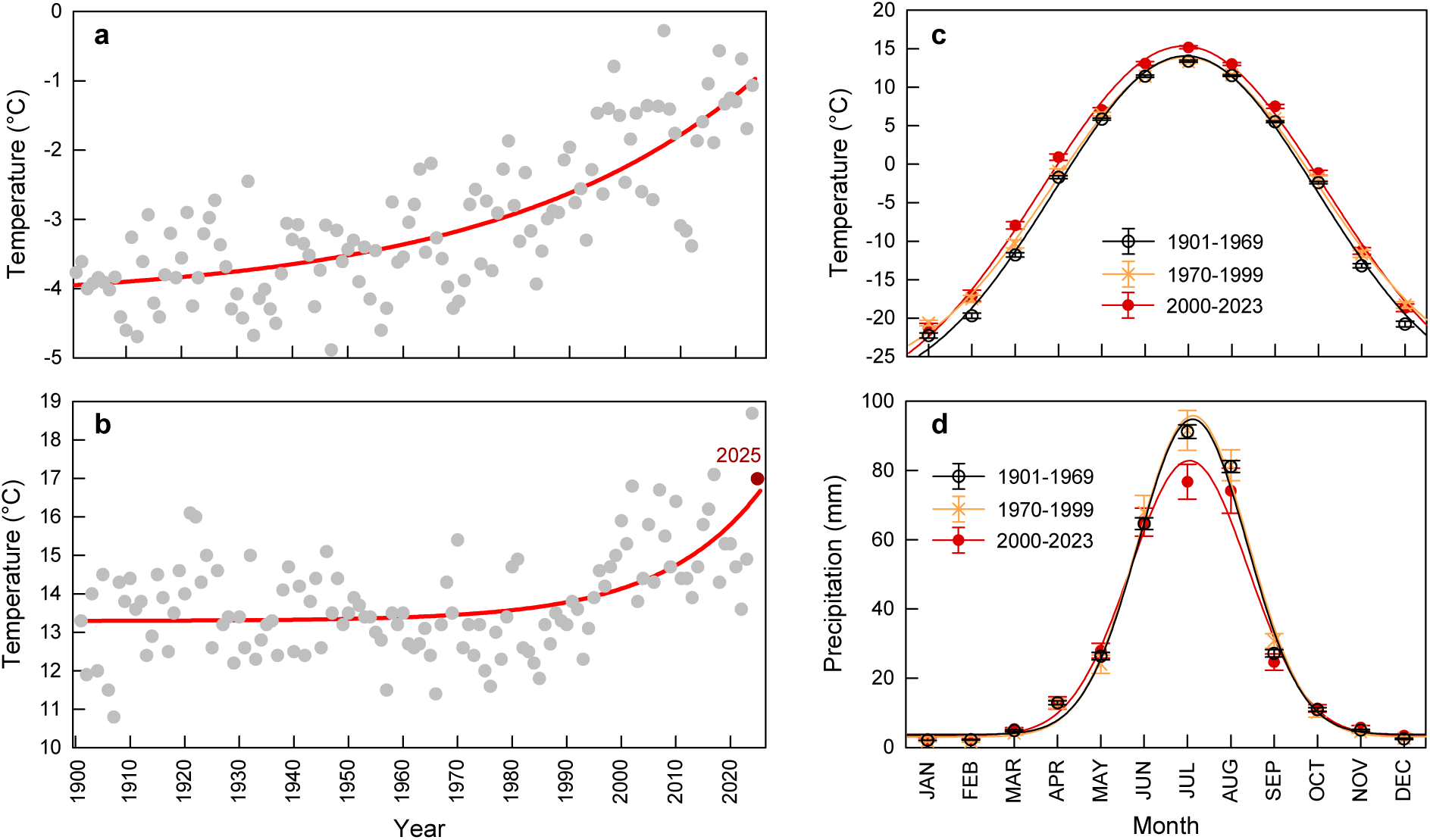
Climate trends since 1901 in the study region modeled from the CRU TS 4.04 dataset edited by the Climate Research Unit of the University of East Anglia, Norwich and the Met Office in Exeter, UK (https://climexp.knmi.nl) at a resolution of 0.5° × 0.5° for the grid field of 48.5 – 49.0 °N and 100.5 – 101.0 °E: (a, b) Trends of (a) mean annual temperature and (b) July temperature; (c) mean monthly precipitation and (d) mean monthly temperatures for three different time intervals (1901–1969; 1970–1999; 2000–2023). For (b) data of July temperatures from 2024 and the year of field work in 2025 (marked in red) were supplemented from the Copernicus ERA5 dataset (https://cds.climate.copernicus.eu).

Mean annual precipitation (MAP) has not changed since 1901 and amounted to 328 mm (lower and upper 5% confidence intervals: 320 and 336 mm). Though MAP remained unchanged since more than a century, there has been a recent reduction of precipitation during the marked peak of rainfall in July (Fig. 1d), which coincides with the annual temperature maximum (Fig. 1c).

### 2.3. Selection of sample trees

Sampling was done in the scope of an established nitrogen fertilization experiment in monospecific *Larix sibirica* Ledeb. forests (Boreal Forest Long-Term Nitrogen Fertilization Experiment, Boreal-LTNE) and was limited to five unfertilized control plots at elevation of 1730–1840 m a.s.l. The experimental setup consisted of five research sites that were distributed over north-facing mountain slopes in a rectangular research area of 15 km in latitude × 5 km in longitude at a mean distance between study sites of 8.7±1.9 km. In each of the five studied forest stands, two dominant to co-dominant *L. sibirica* trees were selected and examined monthly throughout the summer 2025 in June, July, and August. The sampling period covered most of the short growing season in this part of the boreal zone, with bud break between mid and late May and the onset of autumn coloration in *L. sibirica* in September. To explore the variability of heat tolerance between tree individuals, the number of sample trees was expanded to 5 trees in one of the forest stands in June. Field work was conducted on unprotected public forest land and thus did not require formal permission by authorities.

### 2.4. Heat treatments

Branches of the sample trees were collected every month from the sun crown using a Notch Big Shot slingshot (Notch Equipment, Greensboro, North Carolina, USA), which permitted the access to branches of the complete height of the *L. sibirica* trees. Branches were stored in water immediately in the field and kept well-watered under half-shady conditions before heat treatments.

Green needles without visual damage were exposed to heat of defined temperatures (35°C, 40°C, 45°C, 50°C) and incubation times (15, 30, 45, 60, 120, 180, 240 min) in a preheated water bath using immersion boiling devices (Sous Vide Stick Enfinigy, Zwilling, Solingen, Germany). In addition, two types of control samples were analyzed: (1) needles that were not exposed to any heat treatment, and (2) needles that were incubated at 25°C for 4 h to exclude an effect of the incubation treatment that was not caused by heat. These controls were run for all tree individuals and repeated in the three months with the same replication as the heat-treated samples. Needles were first put in paper bags and then sealed in watertight heat-stable plastic bags. The packing in paper bags was done in order to incubate the samples with some air to avoid anaerobic conditions during the heat treatment. The temperature in the sample bags in this experimental setup reliably represents the preset temperature in the water bath, as tested with temperature sensors by Hauck et al. (2025). Incubation and measurements were done with complete needles.

### 2.5. Chlorophyll fluorescence analysis

Ten *L. sibirica* needle per combination of heat/incubation time treatment were analyzed per sample tree and month, resulting in a total of more than 9600 measurements. After the heat treatment, samples were kept in the paper bags and stored in the dark for a minimum of 20 min. Chlorophyll *a* fluorescence analysis was conducted using a Licor Li-600N Porometer/Fluorometer (Licor, Lincoln, Nebraska, USA). The minimal level of chlorophyll fluorescence (*F*_0_) was measured using a weak measuring beam that was not sufficient to induce electron transport at PS II. This was followed by the measurement of maximum fluorescence (*F*_m_) after the application of a saturation pulse (Murchie & Lawson 2013). From the *F*_0_ and *F*_m_ records, we calculated the maximum quantum yield of PS II (*F*_v_/*F*_m_) using the formula *F*_v_/*F*_m_ = (*F*_m_ – *F*_0_)*/F*_m_. The difference in *F*_v_/*F*_m_ of the control samples incubated at 25°C for 240 min and the other control samples that were measured without any heat treatment amounted to 0.013±0.001 and was not significant (Wilcoxon rank sum test, *P*=0.27). Mean *F*_v_/*F*_m_ (±SE) was 0.76±0.01 without any incubation treatment and 0.74±0.01 in the samples exposed to 25°C for 4 h.

### 2.6. Calculation of heat tolerance indices

There are different indices of heat tolerance in plants, all of which respond heavily to incubation time (Neuner & Buchner 2023; Didion-Gency et al. 2025; Hauck & Dulamsuren 2026). Therefore, indices were calculated in dependence of time. We primarily relied on the temperature (*T*_IP_) and *F*_v_/*F*_m_ (*F*_IP_) at the inflection point of the sigmoid *F*_v_/*F*_m_-vs.-temperature relationship at different incubation times, because the inflection point marks the curve section of the most rapid decline in *F*_v_/*F*_m_ and can thus be assumed to reflect severe damage, regardless whether the exact mechanism of this damage at the corresponding temperature is known or not (Hauck & Dulamsuren 2026). We also calculated the temperature of 50% decrease in *F*_v_/*F*_m_ (Krause et al. 2010), though this index (absolute *T*_50_) is not directly supported by a mechanism and can occur under weak or strong reductions of *F*_v_/*F*_m_ values that are likely to be reversible or not (Hauck & Dulamsuren 2026). Furthermore, we calculated *T*_50_’, which is the relative version of *T*_50_, where *F*_v_/*F*_m_ is standardized from 0–100% (Hauck & Dulamsuren 2026). Because *T*_50_’ is also based on the inflection point of the regression line, it represents like *T*_IP_ the temperature range of the most rapid *F*_v_/*F*_m_ decline, but it provides no information on the severity of the decline. Moreover, the standardization results in shifts of *T*_50_’ relative to *T*_IP_ and *T*_50_ (Hauck & Dulamsuren 2026). Even though we therefore preferred the combined specification of *T*_IP_ and *F*_IP_ over *T*_50_ and *T*_50_’, we also calculated these two parameters and present the results in the Supplementary Information, because *T*_IP_ and *F*_IP_ were proposed as heat tolerance indices just recently (Hauck & Dulamsuren 2026).

In addition, we calculated the so-called critical temperature (*T*_crit_), which was defined as the temperature of 50% decrease in *F*_v_/*F*_m_ (Perez & Feeley 2020) and represents the temperature of first signs of membrane damage, which can be assumed to be largely reversible (Kreslavski et al. 2008; Didion-Gency et al. 2025). The calculation of *T*_50_ and *T*_crit_ was based on *F*_v_/*F*_m_ values of dark-adapted untreated control samples, which amounted to 0.76±0.05 in June, 0.78±0.02 in July, and 0.75±0.04 in August. The sigmoid regression equations were solved to *x* (= temperature) using R 4.4.1, when *y* (= *F*_v_/*F*_m_) was set to 50% (*T*_50_) and 15% (*T*_crit_) of the above control values. *T*_IP_ and *F*_IP_ were calculated with the R package ‘nplr’ 0.1-8 under R 4.4.1 by fitting the data to a *N*-parameter model of the generalized logistic function (Richards’ function), starting with 5 parameters. The coordinates of the inflection point of the regression curve, which are included in the ‘nplr’ output, represent *T*_IP_ (*x*-value) and *F*_IP_ (*y*-value). *T*_50_’ corresponds to IC50 in the ‘nplr’ output.

### 2.7. Measurement of microclimate

Microclimate at all 5 study sites was measured with HOBO U23 ProV2 temperature/relative humidity data loggers (Onset Computer Cooperation, Bourne, Massachusetts, USA). These loggers were installed on the surface of *L. sibirica* stems at a height of 1.5 m and at the north side to avoid direct insolation. A total of 12 loggers was distributed through the forests in each of the five studied forest sites (i.e. 60 loggers in total) and temperature values recorded with these loggers were averaged per measurement time for analysis. Measurements were recorded at every full hour. These records were done from August 12, 2024 to August 11, 2025.

### 2.8. Thermal imaging and canopy temperatures

Thermal imaging with a Mavic 3T thermal imaging drone (DJI, Shenzhen, China) was applied to assess canopy temperatures of the *L. sibirica* stands where the branches for experimental heat treatments were collected in parallel to the heat tolerance measurements in the laboratory. Drone flights were conducted in the hot noon and afternoon hours of warm days with clear sky to capture maximum canopy temperatures, which are most relevant for heat stress. Canopy temperatures were measured on automated flights along preset parallel flight routes. Flights were conducted at an altitude of 500 m and thermal and optical images were taken every 1 s, resulting in overlapping image series with a total of 985±63 thermal images per flight.

For determining canopy temperatures, we used the DJI Thermal Analysis Tool 3. Calculating mean values of temperature from entire images easily results in overestimation of canopy temperatures in the not very dense *L. sibirica* forests of Mongolia, because sun-exposed parts of tree trunks, deadwood, open soil and rocks in canopy gaps can attain much higher temperatures than the canopy (Still et al. 2021). Therefore, we refrained from an automated analysis to make sure that temperature maxima in our data reliably reflect *L. sibirica* canopies and not higher temperatures from the ground or deadwood. To achieve this goal, we visually compared optical and thermal images and restricted our temperature readings to image sections that could be completely assigned to *L. sibirica* tree crowns. This manual procedure was preferred for image analysis, as avoidance of overestimation of canopy temperatures was critical for comparing heat thresholds identified during our laboratory measurements with canopy temperatures from the field on warm days. For obtaining a manageable and unbiased sample size, we analyzed every tenth image of each drone flight until 25 images were analyzed. With this random procedure for image selection, we analyzed a total of 375 thermal images, which represents 2.5% of the total images. This is certainly a limitation, but erroneously including higher temperature values from the ground would have led to wrong conclusions in the comparison of canopy temperatures and critical temperatures for PS II damage.

### 2.9. Calculation of thermal safety margins

Thermal safety margins (TSM) were calculated for the differences of ambient peak temperatures to *T*_IP_ and *T*_crit_ (O’Sullivan et al. 2017; Endris & Rehm 2025). Since the TSM represents the difference between two temperatures (in °C), we specify it in K. As peak temperatures, we used (1) the highest air temperatures during our study period, (2) the peak air temperature of 33.0 °C (in 2002) extracted from the ERA5 reanalysis data for 2002, (3) the peak air temperature of the nearest weather station in Erdenemandal of 35.6 °C (in 2002), and (4) the mean and maximum canopy temperatures recorded with thermal imaging from our sample plots on selected days, which represented warm summer days, but not necessarily the hottest days of the growing season. Using linear regression, we could convert standard air temperature in open land at 2 m height to below-canopy temperatures at 1.5 m height, which was recorded during our study, and vice versa. Below-canopy temperatures were converted into estimated of canopy temperatures using linear regression to capture heat maxima of canopy temperatures during periods without direct thermal imaging data.

For the linear regression of temperature time series, we calculated robust Theil-Sen regressions with the R package ‘RobustLinearReg’ 1.2.0. We included daily data of mean and maximum temperatures from 1 JUN to 11 AUG 2025 in our analyses, because heat tolerance measurements where finished by this date. Daily means of below-canopy temperatures could be modeled from ERA5 open land temperatures by Theil-Sen regression (*y* = 0.842 *x* + 1.304; *r*_TS_ = 0.76, *P* < 0.001). Furthermore, maximum below-canopy temperatures were calculated using Theil-Sen regression (*y* = 0.847 *x* + 3.399; *r*_TS_ = 0.55, *P* < 0.001). The goodness of the fit for modeling daily maxima of below-canopy temperatures from open land temperatures was lower than for daily mean temperatures due to positive outliers in the below-canopy data (Fig. S1), resulting in moderate underestimation of below-canopy temperatures if deduced from the ERA5 open land temperatures. Temperature differences between canopy temperatures determined from thermal imaging and below-canopy temperatures were added to measured and modeled below-canopy temperatures to estimate canopy temperatures for the calculation of TSM. These modeled data canopy data and the corresponding ERA5 source data, which were used as input variables in the regression are compiled in Table S1.

We differentiated between variants of the TSM value. The first one (TSM_max_) was calculated from the difference of *T*_IP_ and *T*_crit_ minus the maximum canopy temperature. The second one (TSM_mean_) was calculated from mean maximum canopy temperatures.

### 2.10. Statistical analyses

Arithmetic means are given ± standard errors (SE) for the experimental data of ecophysiological measurements and ± standard deviation (SD) for climate data. Statistical analyses were computed in R 4.4.1. Generalized linear mixed-effects modeling (GLMM) was done with R package ‘lme4’ 1.1-35.5. As *F*_v_/*F*_m_ vs. temperature data follow a sigmoid function, we selected a binomial distribution combined with a logit link function. All numeric predictors were rescaled before analysis in GLMM by subtracted the sample mean from the individual values and dividing the result by the sample’s standard deviation using the ‘scale’ function in R. The sampling month (June to August) was used as numeric parameter from 6–8 and rescaled. The significance of differences between individual sample trees that was tested with an extended sample in one month on one site was also tested with GLMM, where “sample tree” was included as a random factor (Table S2). Predictors were checked for multicollinearity with the R package ‘performance’ 0.12.4. All predictors included in the models had variance inflation factors between 1.0 and 2.1. The interaction effects of temperature × incubation time and time × month (the latter being important for assessing the occurrence of heat acclimation) had to be calculated in two separate GLMM. Generalized linear models (GLM) assuming a Gaussian distribution and applying the inverse link function was employed to test the effect of month and incubation time (after rescaling) on heat tolerance indices (i.e. *T*_IP_, *F*_IP_, *T*_50_, *T*_50_’, and *T*_crit_). The interaction of month × incubation time was only considered for *T*_crit_, as it had no significant effect in the other cases.

Before conducting pairwise comparisons of mean values, we tested for normality with the Shapiro-Wilk test. Based on these tests, we calculated the Wilcoxon rank sum test (*U*-test) for pairwise comparison of not-normally distributed data, Dunn’s test after a Kruskal-Wallis test for multiple comparison of not-normally distributed data, and Tukey’s HSD test for multiple comparison of normally distributed data. Dunn’s test was calculated with the R package ‘dunn.test’ 1.3.6.

## 3. Results

### 3.1. Effects of temperature and incubation time on F_v_/F_m_

Heat exposure in the laboratory caused an increasing reduction of the maximum quantum yield of PS II (*F*_v_/*F*_m_) with both increasing temperature and increasing incubation time (Table 1; Fig. 2). Though both effects were highly significant (*P*<0.001), the regression estimates revealed that the influence of temperature was clearly higher than that of incubation time (Table 1). In addition, there was a negative interaction effect of temperature × incubation time, indicating that long exposure to high temperatures reduced *F*_v_/*F*_m_ particularly strongly (Table 1). Site and sample tree nested within site that were chosen as random factors had no effect on the model results. *F*_v_/*F*_m_ decreased linearly with time at 35°C and 40°C, but exponentially at 45°C and 50°C (Fig. 2). At least after 4 h of exposure to 45°C or 50°C, *F*_v_/*F*_m_ was reduced to 0.000 or close to 0.000 (Fig. 2).

**Fig. 2.**
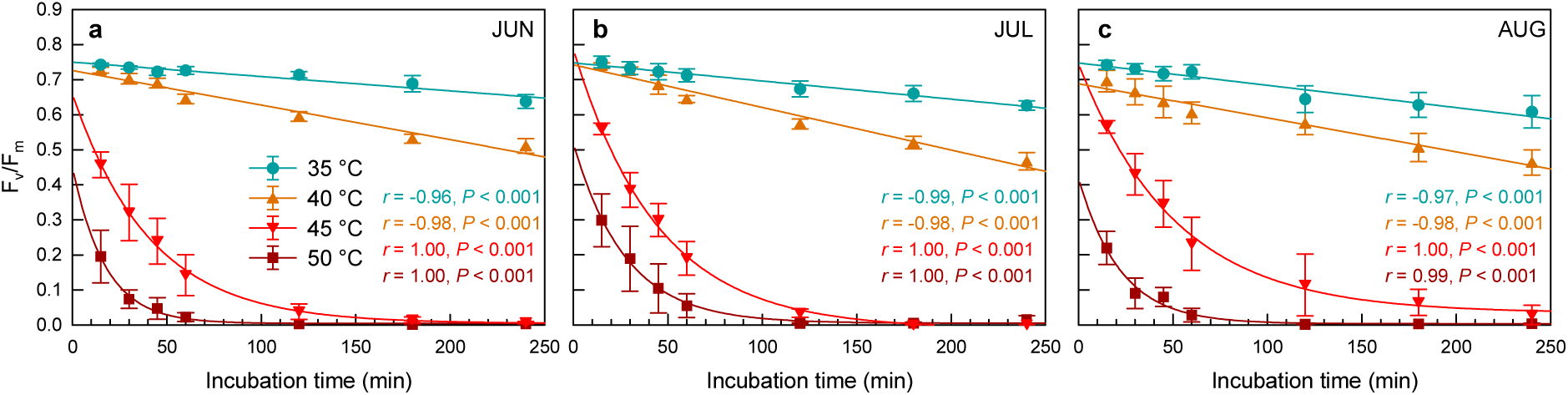
Maximum quantum yield of photosystem II (*F*_v_/*F*_m_) of needles of *Larix sibirica* incubated at 35 – 50°C for up to 4 h in (a) June, (b) July, and (c) August 2025.

**Table 1.**
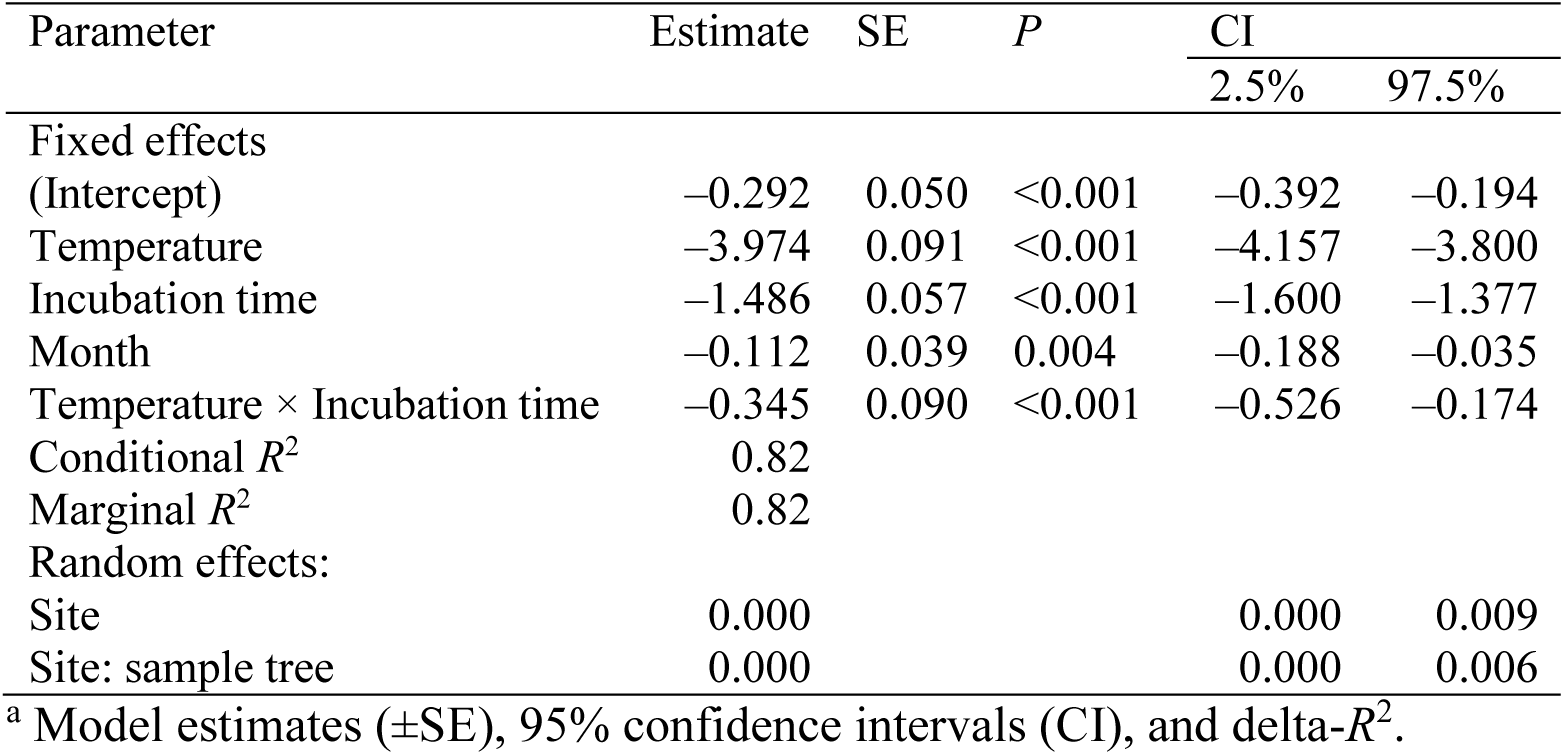
Generalized linear mixed-effects model analyzing effects of temperature, incubation time, month, and the interaction of temperature and incubation time as fixed factors and site and sample tree nested in site (site: sample tree) as random factors on the chlorophyll fluorescence yield of dark-adapted *Larix sibirica* needles^a^.

| Parameter | Estimate | SE | *P* | CI |  |
| --- | --- | --- | --- | --- | --- |
|  |  |  |  | 2.5% | 97.5% |
| Fixed effects |  |  |  |  |  |
| (Intercept) | −0.292 | 0.050 | <0.001 | −0.392 | −0.194 |
| Temperature | −3.974 | 0.091 | <0.001 | −4.157 | −3.800 |
| Incubation time | −1.486 | 0.057 | <0.001 | −1.600 | −1.377 |
| Month | −0.112 | 0.039 | 0.004 | −0.188 | −0.035 |
| Temperature × Incubation time | −0.345 | 0.090 | <0.001 | −0.526 | −0.174 |
| Conditional *R*<sup>2</sup> | 0.82 |  |  |  |  |
| Marginal *R*<sup>2</sup> | 0.82 |  |  |  |  |
| Random effects: |  |  |  |  |  |
| Site | 0.000 |  |  | 0.000 | 0.009 |
| Site: sample tree | 0.000 |  |  | 0.000 | 0.006 |

<sup>a</sup> Model estimates ( ± SE), 95% confidence intervals (CI), and delta-*R*<sup>2</sup>.

### 3.2. Variability between sample trees during heat treatments

Linear mixed-effect modeling did not confirm the occurrence of significant differences between tree individuals of the same forest stand in the response to heat in our experiments (Table S2). Including sample tree identity in the model as a random factor showed no significant effect on the model results, as the conditional *R*^2^ (0.84) and the marginal *R*^2^ (0.83) were nearly identical (Table S2).

### 3.3. Heat acclimation

Even though temperature and incubation time exerted the strongest effects on *F*_v_/*F*_m_ in the experiments, the linear regression model also exhibited weaker, but significant effects of the month of sampling alone and in interaction with temperature (Table 2). While the month of sampling alone reduced the model estimate by –0.10, the interaction of temperature and month increased the estimate by 0.65, indicating an increase in *F*_v_/*F*_m_ during the growing season at high temperatures (Table 2). These GLMM results are reflected by the month-wise comparison of regression curves in Fig. 3. At the lower temperature treatments of 35 and 40°C, where the heat-induced decline of *F*_v_/*F*_m_ was moderate, there was a slight decrease of *F*_v_/*F*_m_ values from June to August (Fig. 3a, b). At the more critical temperature of 45°C, where strong reductions of *F*_v_/*F*_m_ occurred, PS II tolerance to heat increased from June to August (Fig. 3c). In August, *F*_v_/*F*_m_ was consistently higher than that in June across all tested incubation times, which was supported by the results of Dunn’s test (Table S3). At 50°C, differences of the sampling month were only effective at incubation times ≤1 h, because *F*_v_/*F*_m_ was reduced to 0.000 or close to 0.000 after long heat exposure at this temperature (Fig. 3b). At 50°C, *F*_v_/*F*_m_ in both July and August was significantly higher than in June (Table S3).

**Fig. 3.**
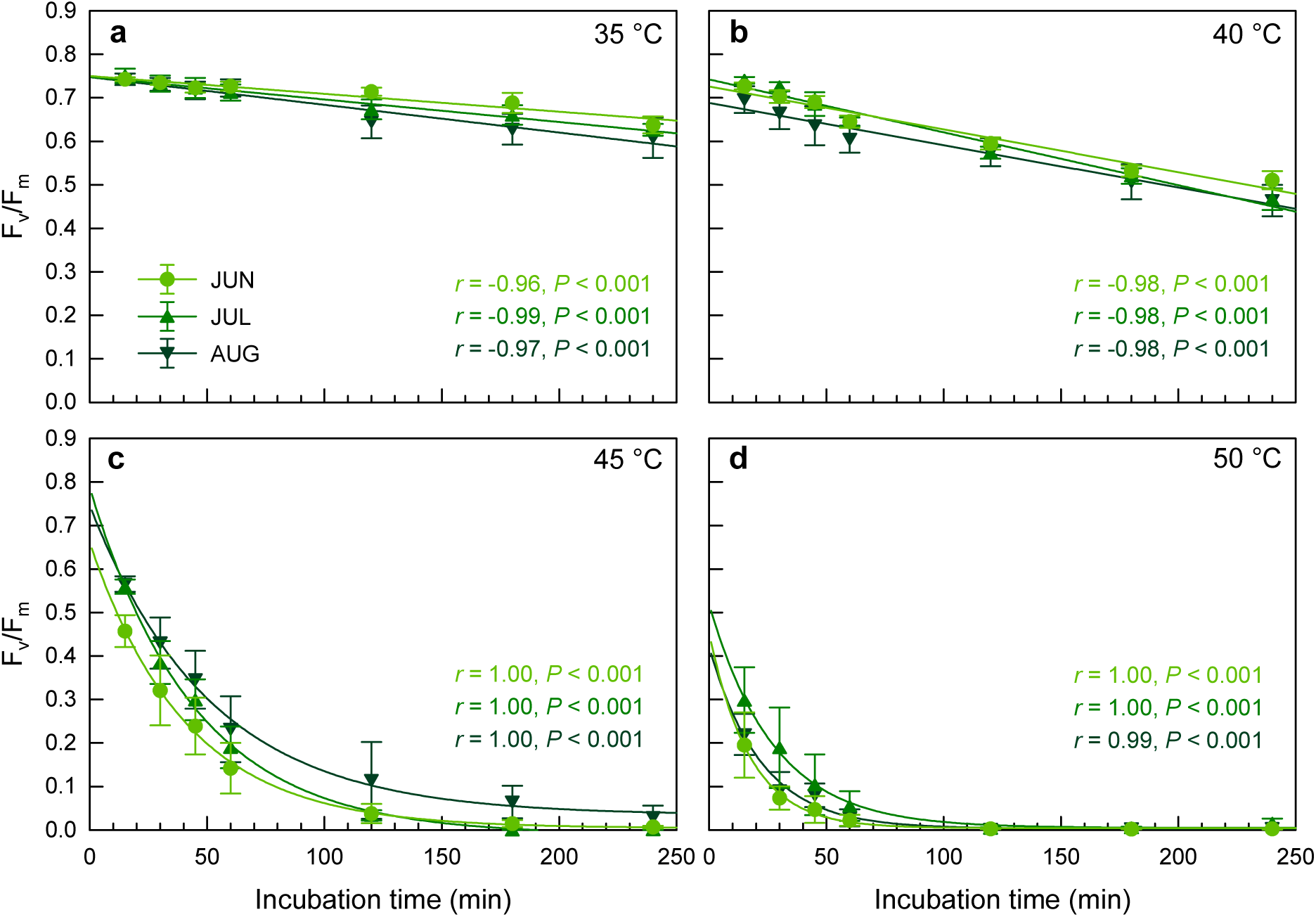
Visualization of potential heat acclimation by comparing *F*_v_/*F*_m_ of needles of *Larix sibirica* incubated in June, July, and August 2025 for up to 4 h at temperatures of (a) 35°C, (b) 40°C, (c) 45°C, and (d) 50°C.

**Table 2.** Generalized linear mixed-effects model analyzing effects of temperature, incubation time, month, and the interaction of temperature and month as fixed factors and site and sample tree nested in site (site: sample tree) as random factors on the chlorophyll fluorescence yield of dark-adapted *Larix sibirica* needles^a^.

| Parameter | Estimate | SE | *P* | CI |  |
| --- | --- | --- | --- | --- | --- |
|  |  |  |  | 2.5% | 97.5% |
| Fixed effects |  |  |  |  |  |
| (Intercept) | −0.150 | 0.040 | <0.001 | −0.228 | −0.072 |
| Temperature | −4.090 | 0.094 | <0.001 | −4.278 | −3.910 |
| Incubation time | −1.430 | 0.051 | <0.001 | −1.532 | −1.331 |
| Month | −0.103 | 0.039 | 0.009 | −0.180 | −0.026 |
| Temperature × Month | 0.647 | 0.069 | <0.001 | −0.514 | −0.782 |
| Conditional *R*<sup>2</sup> | 0.83 |  |  |  |  |
| Marginal *R*<sup>2</sup> | 0.83 |  |  |  |  |
| Random effects: |  |  |  |  |  |
| Site | 0.000 |  |  | 0.000 | 0.009 |
| Site: sample tree | 0.000 |  |  | 0.000 | 0.006 |

<sup>a</sup> Model estimates ( ± SE), 95% confidence intervals (CI), and delta-*R*<sup>2</sup>.

### 3.4. Heat tolerance indices

In agreement with the GLMM results (Tables 1, 2), Fig. 4 showed a strong effect of incubation time and a weaker effect of the month of sampling, when *F*_v_/*F*_m_ was taken up versus temperature. Across all month and incubation times, mean *T*_IP_ was 43.5±0.5°C at an *F*_IP_ of 0.360±0.015 (Table S4). Mean *T*_50_ (43.1±0.4°C) was slightly lower than *T*_IP_, whereas mean *T*_50_’ amounted to 42.1±0.4°C and thus underestimated thermotolerance compared to the two other parameters. Mean *T*_crit_ amounted to 39.8±0.5°C (Table S4). *T*_50_’ was lower than *T*_IP_ and *T*_50_ under almost all combinations of months and incubations times (Fig. S2). *T*_50_ underestimated heat tolerance compared to *T*_IP_ primarily after long heat exposure of 4 h and, in the August data, also at short incubation times (Fig. S2).

**Fig. 4.**
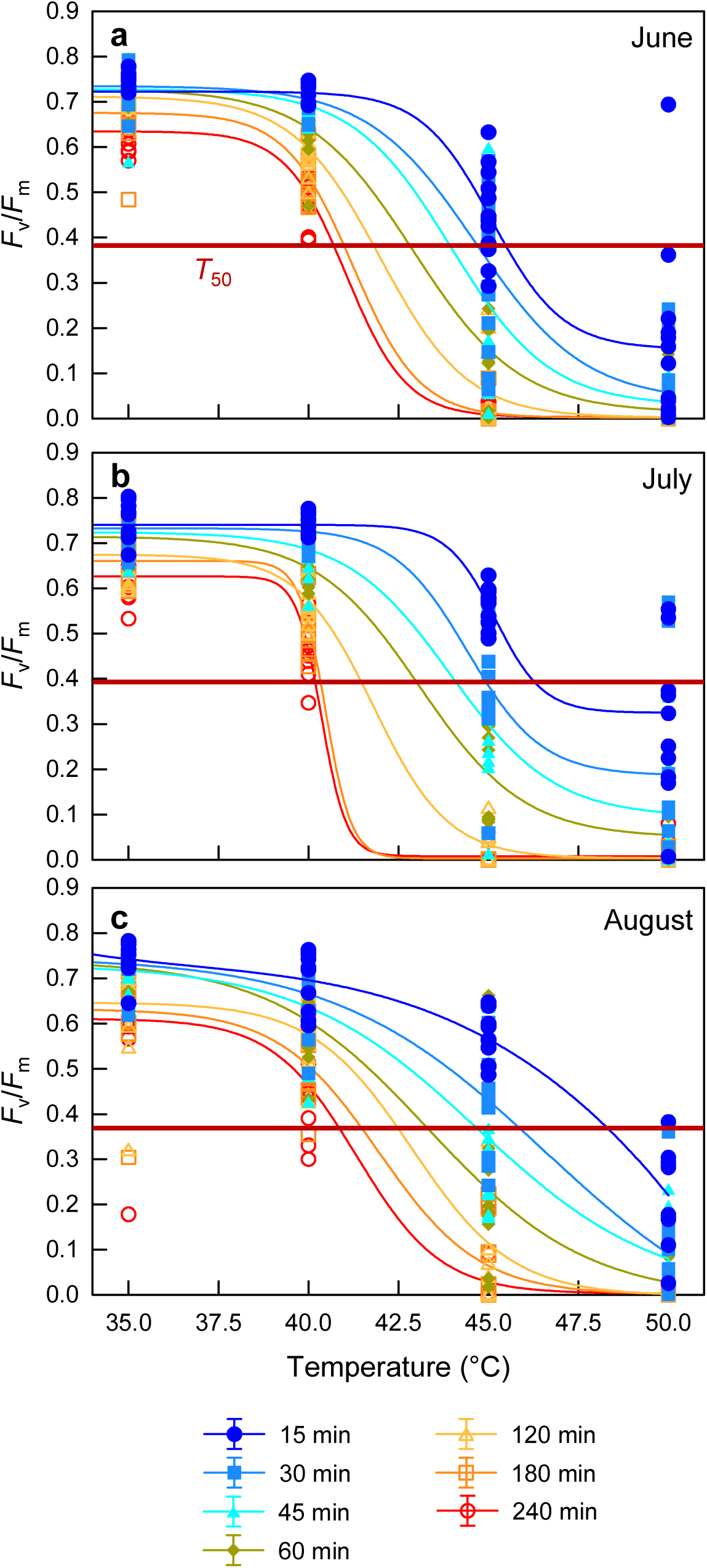
Maximum efficiency at PS II (*F*_v_/*F*_m_) in *Larix sibirica* after heat treatments at 35, 40, 45, and 50°C for 15 to 240 min in different tree species. The red horizontal lines mark the level of 50% of maximum *F*_v_/*F*_m_ (*T*_50_): (a) June, (b) July, (c) August 2025.

After 4 h of heat exposure, mean values from June to August amounted to 41.2±0.0°C for *T*_IP_ and 37.4±0.5°C for *T*_crit_ (Table S4). For samples that were only exposed to heat for 15 min, *T*_IP_ was 47.2±1.6°C and *T*_crit_ was 44.0±0.4°C (Table S4). The corresponding *F*_IP_ values were much lower after 4 h (0.285±0.018) than after 15 min (0.468±0.023) incubation. *T*_50_ amounted to 45.9±0.2°C for 15 min and 40.6±0.2°C for 4 h; the corresponding values for *T*_50_’ were 45.6±0.5°C and 39.8±0.1°C (Table S4).

Comparing *T*_IP_, *T*_50_ and *T*_50_’, the results became more differentiated if data were examined for changes between months, because *T*_IP_ and *F*_IP_ (Fig. 5a, b) indicated the significant changes in the heat response of *F*_v_/*F*_m_ (Fig. 3) over the growing season, whereas *T*_50_ and *T*_50_‘ did not (Fig. S3). This became evident from GLM analyzing the influence of month and incubation time on the heat tolerance indices (Table S5). While the month of sampling exerted a significant effect on *T*_IP_ and *F*_IP_, no such effect was observed for *T*_50_ and *T*_50_’ (Table S5). *T*_IP_ increased from June to August, whereas *F*_IP_ decreased (Fig. 5a, b; Table S5). While *F*_IP_ decreased from 0.44– 0.50 after 15 min of heat exposure to 0.25–0.30 after 4h in June and July, it remained constantly at 0.33±0.01 in August (Fig. 5b).

**Fig. 5.**
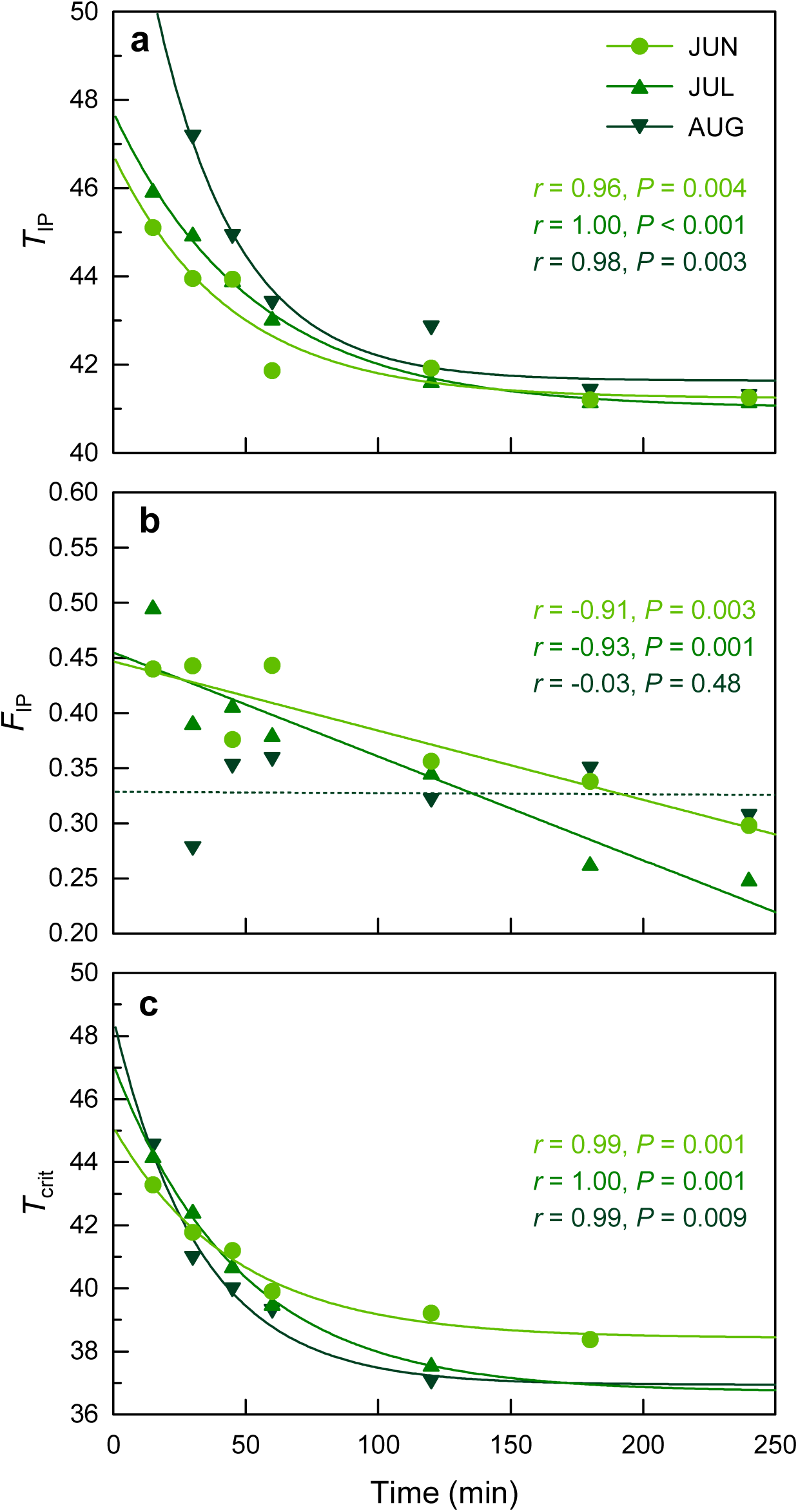
(a) Change of temperature at the inflection point (*T*_IP_) of the regression line for temperature vs. incubation time, (b) *F*_IP_ (i.e. maximum quantum yield [*F*_v_/*F*_m_] at *T*_IP_), and (c) *T*_crit_ (i.e. the temperature of first *F*_v_/*F*_m_ reductions [by 15%]) following different durations of heat exposure from 15 to 240 min from June to August 2025.

*T*_crit_ decreased from June to August (Fig. 5c), indicating increased sensitivity over the growing season, and thus responded the opposite way of *T*_IP_. The differences in *T*_crit_ between months primarily became effective after heat exposure for ≥1 h (Fig. 5c), which is supported by a significant effect of the interaction month × incubation time in addition to an effect of both parameters alone in the GLM (Table S5).

### 3.5. Stand air temperatures at 1.5 m

Air temperature in the *L. sibirica* stands at 1.5 m height showed a wide seasonal amplitude (Fig. 6). Daily mean temperature (±SD) in the 1-year interval from August 2024 to August 2025 was 1.1±12.0°C. The annual means of daily minimum and maximum temperatures amounted to –5.1±11.2°C and 7.6±13.3°C. The highest daily mean temperature value was 20.6°C, while the highest absolute temperature maximum recorded was 30.8°C. The coldest day had a mean temperature of –24.4°C and a minimum temperature of –27.5°C.

**Fig. 6.**
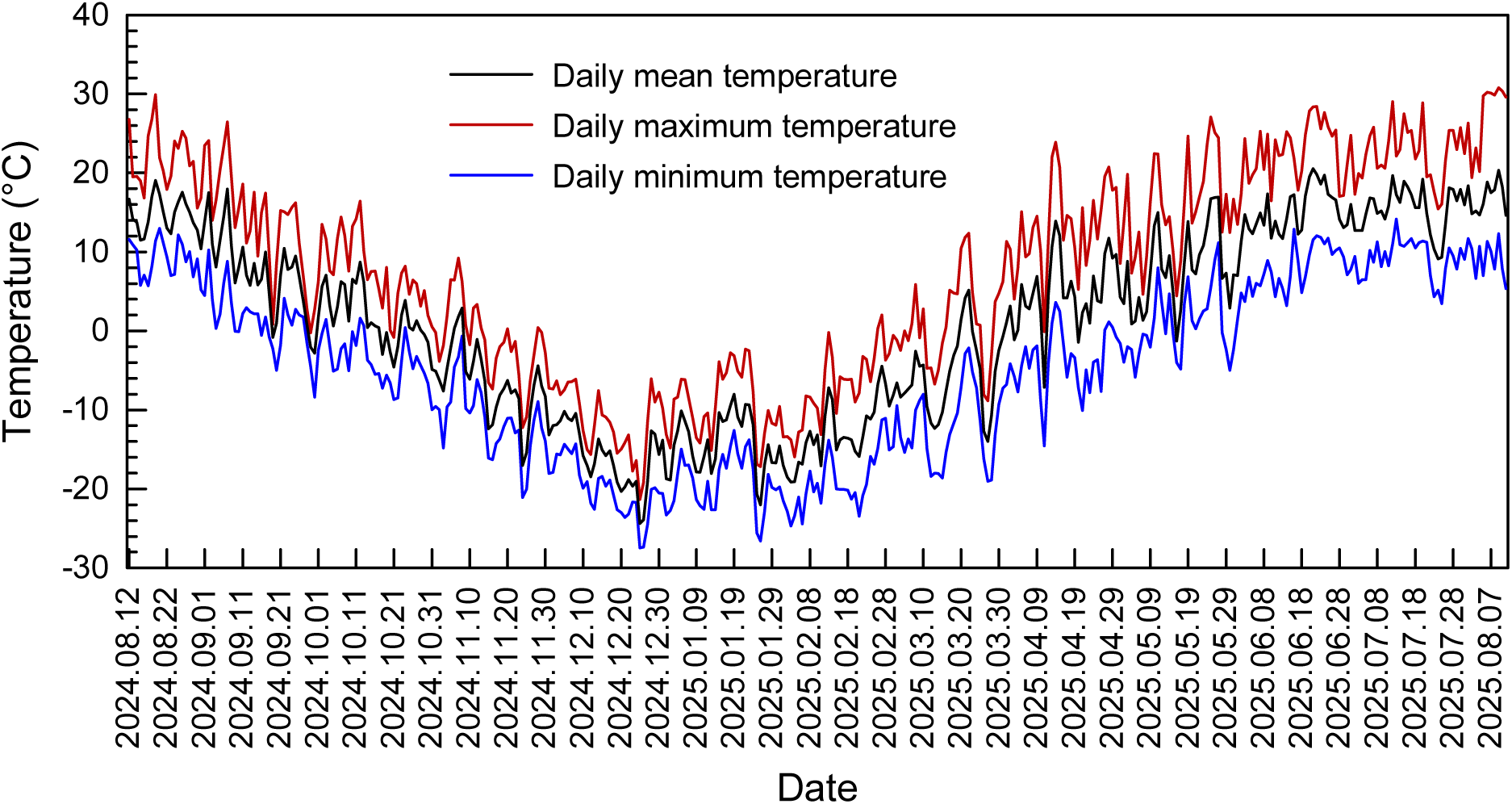
Daily mean, maximum, and minimum air temperatures at 1.5 m height in the studied *Larix sibirica* stands from 12 August 2024 to 11 August 2025.

The absolute temperature maxima of the study months in June, July, and August 2025 (data only available until 11 AUG 2025, when heat experiments were finished) ranged from 28– 31°C, while mean maximum temperatures (±SD) amounted to 22.5±4.0°C (June), 22.5±3.5°C (July), and 27.2±4.3°C (August; Table S6). Mean temperatures ranged from 15–17°C in this period (Table S6).

### 3.6. Canopy temperatures and their relationship to air temperature

Since canopy temperatures were recorded on warm days with clear sky, the instantaneous air temperatures during the drone flights (measured below the canopy at 1.5 m height; Fig. 7a) exceeded the mean air temperatures measured by the same sensor from June to August (Table S6) by more than 10 K. While mean temperature (±SD) from June to August was 15.4±2.9°C (Table S6), the instantaneous below-canopy temperatures during the drone flights were 24.6±2.5°C (Fig. 7a).

**Fig. 7.**
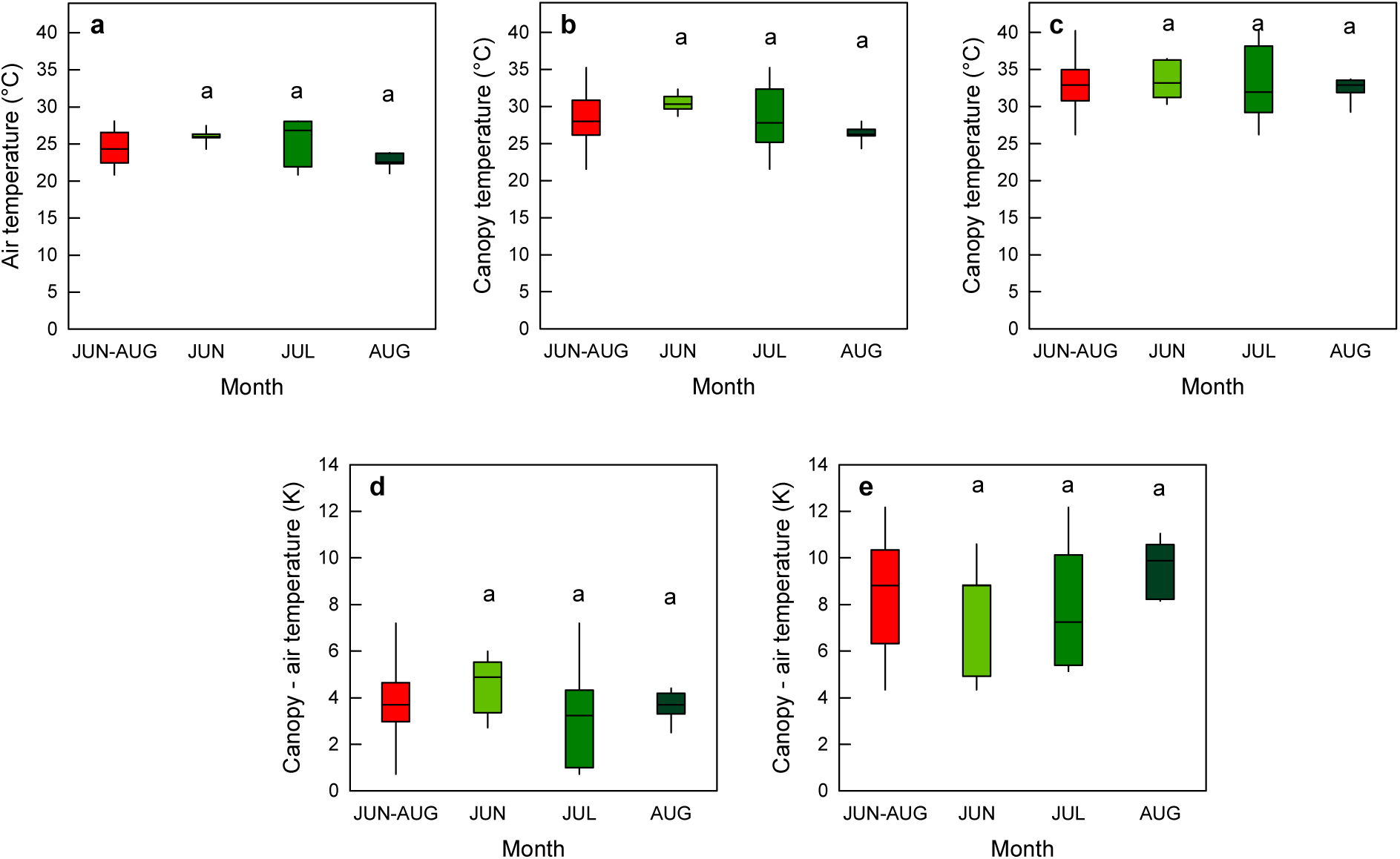
Instantaneous (a) air temperature in the *Larix sibirica* stands at 1.5 m, (b) mean and (c) maximum canopy temperatures, and differences of (d) mean and (e) maximum canopy minus air temperatures minus air temperature at the time of measurement. Boxplots for June to August (red) and individual months. Means of data from individual months sharing the same letter, do not differ significantly (Tukey’s HSD test, *P*≤0.001)

Mean canopy temperatures (±SD) amounted to 30.5±1.4°C in June, 28.4±5.5°C in July, and 26.3±1.3°C in August, with a mean of 28.4±3.6°C for all drone flights from June to August (Fig. 7b). Maximum canopy temperatures recorded in the thermal images exceeded mean canopy temperatures by 4.6±1.6 K. Maximum values were 33.5±2.8°C in June, 33.2±5.9°C in July, 32.3±1.8°C in August, and 33.0±3.7°C as the mean of June to August (Fig. 7c).

The mean difference between mean canopy temperatures and below canopy temperatures at 1.5 m was 3.8±1.7 K (Fig. 7d). These values varied between 4.5±1.4 K, 3.3±2.7 K, and 3.6±0.8 K during the June, July, and August drone flights (Fig. 7d). The maximum canopy temperatures exceeded the instantaneous below-canopy temperatures by 8.4±2.5 K (June to August) and by 7.5±2.7 K (June), 8.0±3.1 K (July), and 9.6±1.3 K (August) in the individual months (Fig. 7e). The difference between mean canopy temperature and the instantaneous air temperature below the canopy increased with increasing air temperature (Fig. S4).

### 3.6. Thermal safety margins

Reference canopy temperatures used for TSM calculation are compiled in Table 3. They included (1) mean (28.4±0.9°C) and maximum (33.0±0.9°C) values (±SE) from thermal imaging from June to August 2025, (2) record temperatures from ERA5 reanalysis from the study area with a mean temperature of 35.2°C and a maximum temperature of 37.9°C modeled for the hottest day of the period 2000–2025 in 2002, and (3) estimated mean (37.4°C) and maximum (42.0°C) canopy temperatures of 2002 from derived from open land data of the nearest instrumental weather station in Erdenemandal, at a location where also *L. sibirica* forests occur. Relevant peak canopy temperatures thus ranged from 33–42°C (Table 3). These temperatures were compared to mean values of *T*_crit_ of 39.8±0.5°C and of *T*_IP_ values of 43.5±0.5°C across all incubation times and to means of these indices for the individual incubation times across the three months of study (Table S4).

**Table 3.**
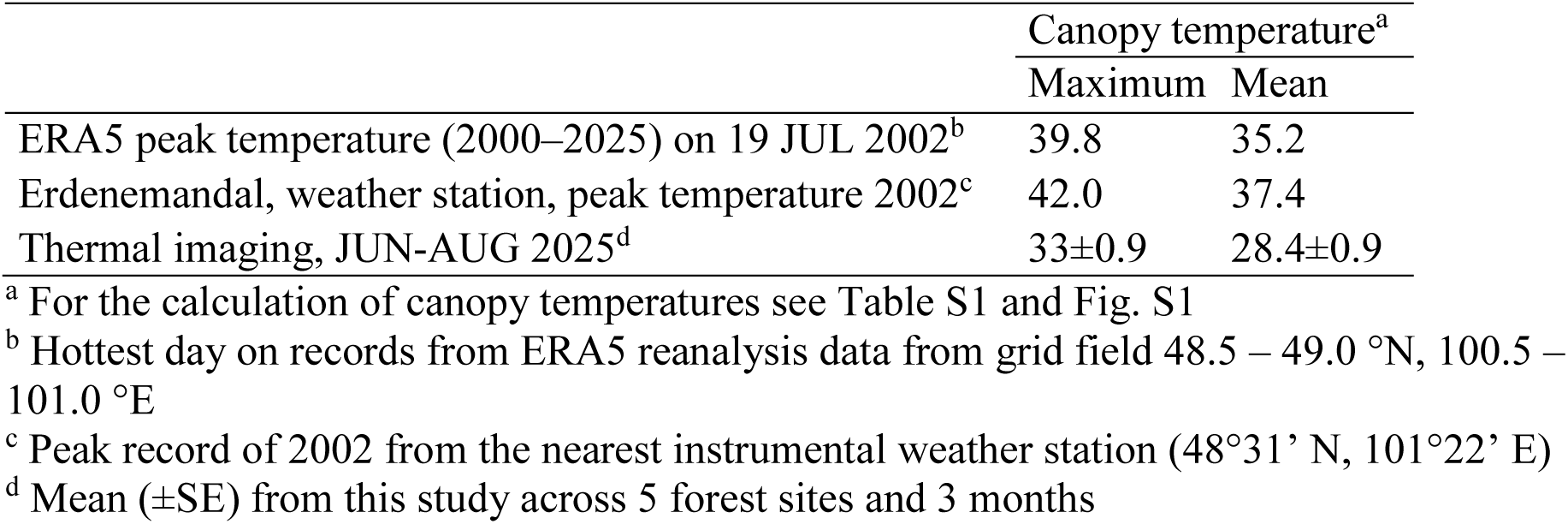
Maximum and mean canopy temperatures used for the calculation of thermal safety margins (TSM).

The canopy temperatures recorded from our sample plots by thermal imaging from June to August 2025 did not reach *T*_crit_ and *T*_IP_ (Fig. 8). TSM_mean_ was as high as 12.2 K relative to *T*_crit_ (Fig. 8d) and 14.7 K for *T*_IP_ (Fig. 8b). Even TSM_max_ was still high with 7.6 K for *T*_crit_ (Fig. 8c) and 10.1 K for *T*_IP_ (Fig. 8a).

**Fig. 8.**
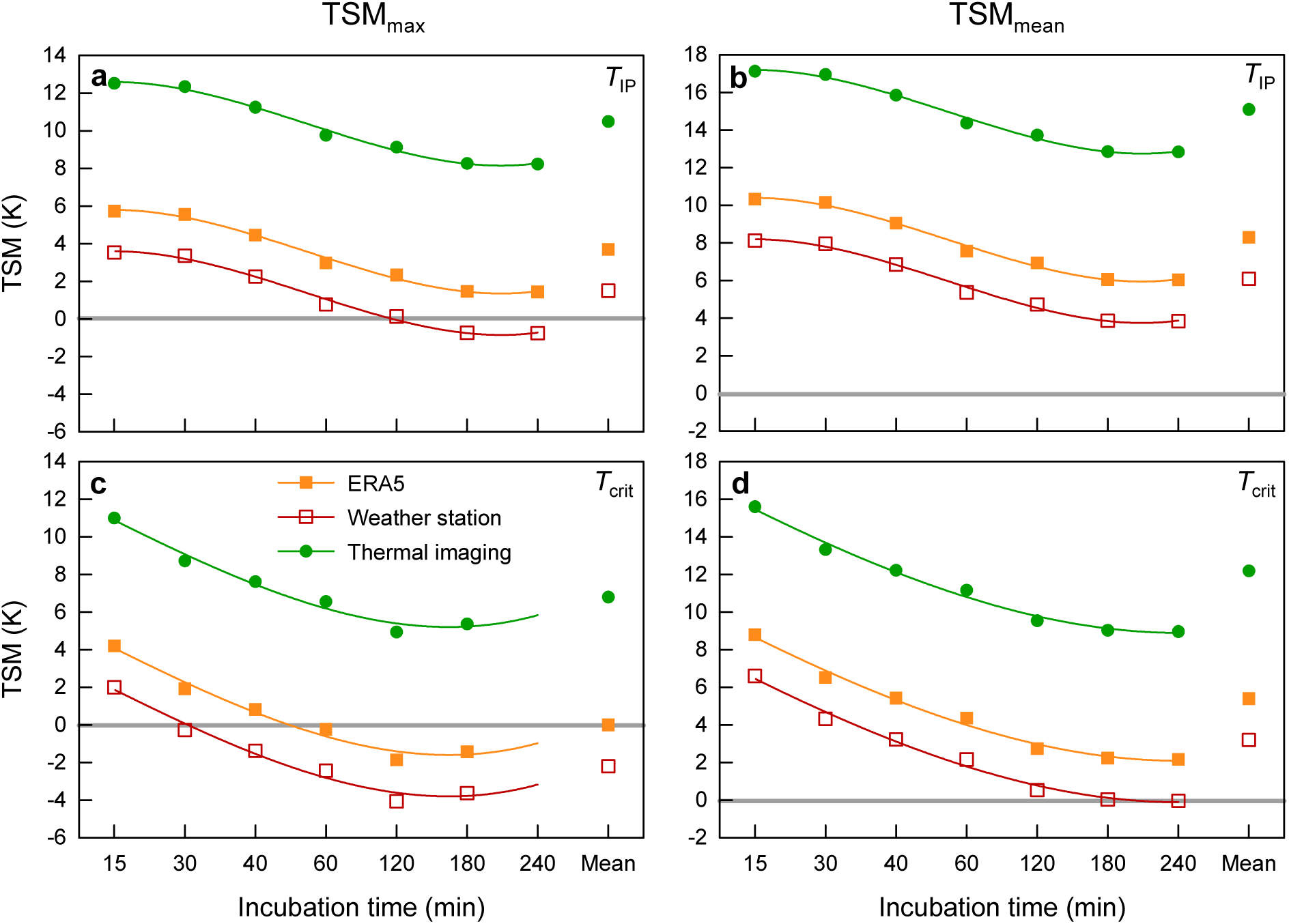
Thermal safety margins (TSM) in *Larix sibirica* based on (a, c) maximum (TSM_max_) and (b, d) mean (TSM_mean_) canopy temperatures calculated relative to (a, b) *T*_IP_ and (c, d) *T*_crit_ values determined after 15 min – 4 h of heat exposure. Mean represents the arithmetic means at all incubation times. Values from thermal imaging are mean values of June to August 2025. ERA5 and weather station represent modeled canopy temperatures during heat extremes of 2002 (hottest heat waves from 2000–2025). These canopy temperatures were modeled from ERA5 reanalysis data for the grid field 48.5 – 49.0 °N, 100.5 – 101.0 °E and weather station data (Erdenemandal, 48°31’ N, 101°22’ E). The grey horizontal line marks the threshold of TSM = 0, when canopy temperatures exceed *T*_IP_ and *T*_crit_.

While the canopy temperatures determined in 2025 with thermal imaging naturally did not cover the entire range of temperatures, the hottest days of the period from 2000–2025 that occurred in 2002 yielded a different result with regard to TSM. TSM_max_ calculated for *T*_crit_ was in the negative range during these heat extremes if heat exposure lasted for ≥30–60 min (Fig. 8c). If calculated with *T*_IP_, TSM_max_ became negative after 3 h of heat exposure (Fig. 8a). TSM_mean_ calculated with *T*_IP_ was positive throughout even during the most extreme heat maxima with values ranging from 3.8–10.2 K (Fig. 8b). Even for *T*_crit_, TSM_mean_ was mostly positive, but was 0.0°C after 3–4 h heat exposure, if calculated for canopy temperatures that were modeled from the data of the nearest weather station (Fig. 8d).

## 4. Discussion

### 4.1. Response to heat treatments

The experimental results revealed a clear influence of both temperature and incubation time, which is consistent with previous studies from temperature broadleaved and coniferous tree species (Neuner & Buchner 2023; Hauck et al. 2025). In contrast to the control samples incubated at 25°C, even temperatures of 35°C and 40°C showed an increasing reduction with increasing duration of the heat treatment and temperature. The nonlinear decrease of the maximum quantum yield of PS II (*F*_v_/*F*_m_) with time at 45°C and 50°C, in contrast to the linear decline at lower temperatures, indicates the existence of a threshold for severe heat damage in PS II between 40°C and 45°C, which is in line with the mean *T*_IP_ of 43.5±0.5°C, with the physiological and biochemical literature dealing with PS II heat tolerance (Allakhverdiev et al. 2008; Mohanty et al. 2012) and with our first hypothesis.

Threshold temperatures for damage from laboratory experiments between different studies have to be compared with care, as the exact temperature thresholds, from which on a certain level of damage is observed, is influenced by the details of the experimental conditions (Didion-Gency et al. 2025; Hauck & Dulamsuren 2026). Regression analysis confirmed that the effect of temperature on *F*_v_/*F*_m_ was clearly higher than that of incubation time, which is plausible, as at high temperatures already short treatments can be assumed to cause damage. Water splitting at the oxygen-evolving complex of PS II is particularly heat-sensitive (Berry & Björkman 1980; Mohanty et al. 2012) and can be assumed to have contributed to the decline of *F*_v_/*F*_m_ already at 35°C, as water splitting was shown to be inhibited already at 32°C in potato leaves (Havaux 1993). Grana stacks of thylakoid membranes, which are associated with PS II (Gu et al. 2022), remained intact in isolated chloroplasts at 35°C, but had completely disintegrated at a temperature of 40°C (Gounaris et al. 1983). Grana destacking is connected with the dissociation of the light-harvesting complex from the core complex of PS II (Gounaris et al. 1984). Heating to temperatures of 45°C and 50°C caused widespread phase separation of non-bilayer lipids in the thylakoid membranes of isolated chloroplasts (Gounaris et al. 1984). Non-bilayer lipids constitute ca. 60% of the chloroplast lipids and have functions in the organization of the thylakoid structure (Garab et al. 2022).

To assess the species-specific heat tolerance of *L. sibirica*, a direct comparison to other tree species can be done with the data of Hauck et al. (2025), where exactly the same methodology was used. This comparison shows that *L. sibirica* is a heat-sensitive species. Especially at the critical temperature of 45°C, when *F*_v_/*F*_m_ starts to decrease exponentially with increasing incubation time (Fig. 2), the decrease of *F*_v_/*F*_m_ is stronger and faster than in the other conifers studied by Hauck et al. (2025), which included *Abies alba*, *Picea abies*, *Pinus sylvestris*, and *Pseudotsuga menziesii*. The difference was even greater compared to ten temperate broadleaved tree species of the genera *Acer*, *Fagus*, *Quercus*, and *Tilia* (Hauck et al. 2025). The generally lower heat tolerance of conifers compared to broadleaved angiosperm trees matches with observations based on short-term heat treatments and on the calculation of *T*_50_’ values for temperate tree species by Kunert & Hajek (2022), Kunert et al. (2022) und Münchinger et al. (2023).

Based on values calculated after 4 h of heat exposure, *T*_IP_ in *L. sibirica* (41.2±0.0°C: mean from June to August; Table S4) was lower compared to values available from *Picea abies* (42.8°C) and *Pinus sylvestris* (42.4°C), slightly lower than in *Abies alba* (41.6°C), but higher than that of *Pseudotsuga menziesii* (40.6°C) (Hauck & Dulamsuren 2026). The temperate-subalpine European *Larix decidua* is less heat-tolerant (*T*_IP_ = 39.8°C; Hauck & Dulamsuren 2026) than *L. sibirica*. This comparison with temperate tree species and with the boreal-temperate *Picea abies* and *Pinus sylvestris* suggests that *L. sibirica* as a species of cold boreal forests in Siberia and northern Central Asia (Mamet et al. 2019) has evolved an only limited heat tolerance, which matches with subzero or near-zero mean annual temperatures at many places of Mongolia’s boreal forest region (Dagvadorj et al. 2009; Hauck et al. 2016), including our study area.

### 4.2. Heat acclimation

Heat acclimation in trees is generally not very well studied and most studies where its occurrence has been suggested did not show a continuous increase in PS II heat tolerance over the growing season. Húdoková et al. (2022) conducted heat treatments of *Quercus petraea* at two points in time in June and August and found higher *F*_v_/*F*_m_ values during the repeat measurements in August, which is suggestive of heat acclimation during summer. *Abies alba*, *Fagus sylvatica* and *Pinus sylvestris* did not show such an increase in heat tolerance from June to August in the same study (Húdoková et al. 2022). *Picea abies* and *Abies alba* had higher *F*_v_/*F*_m_ during summer compared to spring (Petrik et al. 2023; Hauck et al. 2025). Newly formed needles were even less heat-tolerant in spring than older ones in both species (Hauck et al. 2025). A recent study of Hauck et al. (2026) pointed to steady heat acclimation over the growing season in *Fagus orientalis* and *Pseudotsuga menziesii*, but basically not in *Fagus sylvatica*.

The results of the present study suggest that there was limited heat acclimation over summer with respect to severe heat damage of PS II at temperatures of ≥45°C, which thus supports our second hypothesis. This was corroborated from both GLMM results for modeling *F*_v_/*F*_m_ values and GLM results for modeling *T*_IP_ and *F*_IP_. Remarkably, this apparent acclimation to heat to avoid severe PS II dysfunction contrasted with a weakening of the PS II tolerance to moderate heat. This is suggested by decreasing *F*_v_/*F*_m_ values at given combinations of temperature and incubation time under moderate heat of 30–40°C over the summer and, moreover, by decreases in *T*_crit_ after long heat treatments from June to August. These differences were significant in GLMM (*F*_v_/*F*_m_) and GLM (*T*_crit_) as well.

We are not aware of any previous reports of such contrasting responses, in which increased tolerance of PS II to extreme heat is accompanied by a parallel decrease in tolerance to moderate heat. It could be the result of the rearrangement of resources in order to prevent critical, potentially lethal damage at high temperatures. Fortifying membranes and proteins with HSP and changes in the membrane lipid composition (Zheng et al. 2011; Reszczyńska & Hanaka 2020; Zhu et al. 2024) has metabolic costs and *L. sibirica* growing under the ultracontinental climate of Mongolia has only limited resources to spend (Liu et al. 2019; Dulamsuren et al. 2023).

The growing season in Mongolia’s boreal forest region is extremely short from mid to late May to September, of which we covered most with our study from June to August. In addition, *L. sibirica* has like many conifers a sensitive stomatal regulation, which reduces the risk of drought damage, but lowers the photosynthetic carbon gain (Dulamsuren et al. 2009; Dulamsuren & Hauck 2021) and results in slow growth (Dulamsuren et al. 2010; Liu et al. 2013; Khansaritoreh et al. 2017). Moreover, the species’ non-structural carbon (NSC) pools are needed to constitute drought tolerance by osmotic and elastic adjustment (Dulamsuren et al. 2023; Dulamsuren 2026) and herbivore defense by producing secondary metabolites (Subbotina et al. 2025). This might pose the need to limit heat acclimation on improving the tolerance to extreme heat over summer while accepting the possibility of weak reversible damage due to moderate heat. This explanation is speculative at the present state of knowledge, but should be worth scrutinizing in future studies.

### 4.3. Interpretation of heat tolerance indices

From a methodological perspective, our results support the combined use of *T*_IP_ and *F*_IP_ to characterize the severe, probably largely irreversible PS II heat damage (Hauck & Dulamsuren 2026) and of *T*_crit_ for the quantification of initial, but largely reversible PS II heat damage (Perez & Feeley 2020; Didion-Gency et al. 2025). The contrasting (statistically significant) adaptive responses of *T*_IP_ and *T*_crit_ to hot summer weather in *L. sibirica* clearly suggest the use of both metrics. The fact that the changes in heat tolerance over the summer could be identified as a significant effect of the month for *T*_IP_ and *F*_IP_ (and *T*_crit_) in the regression models, but not for *T*_50_ or *T*_50_’, underscores their greater discriminatory power of the former. This higher discriminatory power is attributable to the better mechanistic substantiation of *T*_IP_ than of *T*_50_ and to the lack of fluorescence data transformation to relative values in the calculation process compared to *T*_50_’ (Hauck & Dulamsuren 2026), which apparently makes a difference in the explanatory power, even though the absolute values of *T*_50_ and to a lesser degree of *T*_50_’ may at least partly be similar to *T*_IP_.

In addition, our study points to the need for employing varying and long incubation times in heat experiments that target on the calculation of TSM. Although it is possible that leaves experience heat extremes for few minutes only during lulls under otherwise windy weather conditions (Curtis et al. 2014; Kunert et al. 2026b), typically extreme compound heat and drought events last for longer intervals and intensify over time during calm weather (Tripathy et al. 2023; Ji et al. 2025). On the hottest days on records in 2002, modeled maximum canopy temperatures over 30°C lasted for 7–9 h and occurred during a heat spell that lasted for several days. We found a clear decrease in TSM with increasing duration of heat exposure, leading to very different conclusions depending on whether 15 min or 4 h of heat treatment was applied for determining *T*_IP_ and *T*_crit_, which are included in the calculation of TSM. Endris & Rehm (2025) in their seminal paper on TSM in North American temperate broadleaves only found critically low or negative TSM values if they compared leaf temperatures with *T*_crit_, but not *T*_50_. It is obvious that this result was affected by selecting a short incubation period of 15 min for determining these parameters (Endris & Rehm 2025).

### 4.4. Relevance of physiological thresholds of heat tolerance under field conditions

Our selective measurements of canopy temperatures on warm summer days with clear sky and our 1-year records of stand air temperature suggest that *L. sibirica* is usually not threatened by lethal damage due to heat on north-facing mountain slopes, which represent the species’ main habitat in Mongolia’s boreal forest. This result confirms our third hypothesis. However, our results also show that PS II damage during extreme heat events, like in summer 2002, have the capacity to contribute to increased mortality of *L. sibirica*, as even TSM_max_ calculated with *T*_IP_ became partly negative after heat exposure for more than 2 h, indicating a high probability of irreversible heat damage (Didion-Gency et al. 2025).

In contrast to Slot et al. (2021) and Endris & Rehm (2025), we do not assume that *T*_crit_ is an indicator of critical irreversible PS II damage (Hauck & Dulamsuren 2026), which is consistent with Kreslavski et al. (2008) and Didion-Gency et al. (2025). Therefore, the terminology of *T*_crit_ as the “critical temperature” is somewhat misleading. However, *T*_crit_ is nevertheless an important parameter, because it indicates a temperature range, at which the heat-sensitive CO_2_ assimilation by the Calvin cycle is already substantially reduced (Haldimann & Feller 2005; Perez et al. 2021) and where in addition first decreases in *F*_v_/*F*_m_ indicate the impairment of the light reaction of photosynthesis, but at the same time the need for additional carbon for the repair mechanisms (Kreslavski et al. 2008; Mohanty et al. 2012). If heat is combined with drought, these requirements can probably hardly be met, as *L. sibirica* closes its stomates early under drought (Dulamsuren et al. 2009; Dulamsuren & Hauck 2021) and also invests more carbon than other trees species in generating low osmotic potentials and increasing cell wall rigidity for osmotic and elastic adjustment (Dulamsuren et al. 2023; Dulamsuren 2026).

## 5. Conclusions

Our experimental heat treatments suggest that *L. sibirica* is more sensitive to heat than most temperate coniferous and even more than temperate broadleaved tree species tested for PS II thermotolerance, so far. However, it is more tolerant to heat than the temperate-subalpine *L. decidua* and the temperate *Pseudotsuga menziesii*. The relatively low heat-tolerance of *L. sibirica* is consistent with its distribution in cold boreal forests of Siberia and northern Central Asia, including Mongolia, and explains why the species does not extend to the temperate zone. Whether low heat tolerance is a general trait of boreal forest trees has still to be explored. Our results from drone-borne thermal imaging showed that heat does not pose a lethal risk for *L. sibirica* during average warm summer days with clear sky in the species’ main habitat in Mongolia’s boreal forest, whereas the hottest days on record in the study region since 2000 had the potential to cause heat-induced mortality. Yet, also moderate heat impaired the light reaction of photosynthesis, as evidenced by lowered *F*_v_/*F*_m_. This observation suggests that heat acts as an additional stressor in *L. sibirica* that lowers carbon assimilation, but increases the demand for carbon for repair of heat damage and for heat acclimation. This adds to the challenges that other stress factors, including drought and herbivores, pose to the trees’ carbon budget, all of which are become increasingly significant due to climate change.

## Acknowledgments

We are thankful to the Forest Office (Mr. D. Batbaatar, Mr. B. Tseveldorj) in Jargalant, Arkhangai district, Mongolia for multiple support during field work.

## CRediT authorship contribution statement

**Choimaa Dulamsuren:** Conceptualization, Funding acquisition, Methodology, Investigation, Data curation, Formal analysis, Writing – original draft, Writing – Review & Editing. **Temuulen Abbas Jayamaran:** Investigation, Data curation, Writing – Review & Editing. **Germar Csapek:** Investigation, Writing – Review & Editing. **Erdenechuluun Naranbayar:** Investigation. **Tumendelger Uitumen:** Investigation. **Dugersuren Amarjargal:** Investigation. **Gurbazar Byamba-Yondon:** Investigation. **Davaadorj Saindovdon:** Investigation. **Tungalag Munkhzul:** Investigation. **Ganbaatar Batsaikhan:** Investigation, **Markus Hauck:** Conceptualization, Methodology, Writing – Review & Editing.

## Data availability statement

Data will be made available on Zenodo after acceptance.

---

**Table S1.** Estimation of peak canopy temperatures using the regression equations from Fig. S1 for mean and maximum canopy temperatures on days with record heat during 2000–2025. Data are modeled from ERA5 reanalysis data and weather station data from open land.

|  | Temperature,<br>open land<br>(°C) | Temperature,<br>below<br>canopy (°C) | Canopy temperature (°C) |  |
| --- | --- | --- | --- | --- |
|  |  |  | Mean<sup>a</sup> | Maximum<sup>b</sup> |
| ERA5 peak temperature (2000–2025)<br>on 19 JUL 2002<sup>c</sup> | 33.0 | 31.4 | 35.2 | 39.8 |
| Erdenemandal, weather station, peak<br>temperature 2002<sup>d</sup> | 35.6 | 33.6 | 37.4 | 42.0 |
| Thermal imaging, JUN–AUG 2025<sup>e</sup> | . | . | 28.4±0.9 | 33±0.9 |

<sup>a</sup> Modeled from open land temperature using $y = 0.842x + 1.304$
<sup>b</sup> Modeled from open land temperature using $y = 0.847x + 3.399$
<sup>c</sup> Hottest day on records from ERA5 reanalysis data from grid field 48.5 – 49.0 °N, 100.5 – 101.0 °E
<sup>d</sup> Peak record of 2002 from the nearest instrumental weather station (48°31' N, 101°22' E)
<sup>e</sup> Mean (±SE) from this study across 5 forest sites and 3 months

**Table S2.** Generalized linear mixed-effects model (GLMM) analyzing effects of temperature and incubation time (fixed factor) and sample tree (*N*=5; random factor) on the chlorophyll fluorescence yield of dark-adapted *Larix sibirica* needles at one site in June^a^.

| Parameter | Estimate | SE | *P* | CI |  |
| --- | --- | --- | --- | --- | --- |
|  |  |  |  | 2.5% | 97.5% |
| Fixed effects |  |  |  |  |  |
| (Intercept) | 0.340 | 0.116 | 0.003 | –0.569 | –0.115 |
| Temperature | –5.191 | 0.333 | <0.001 | –5.889 | –4.579 |
| Incubation time | –1.743 | 0.165 | <0.001 | –2.083 | –1.435 |
| Conditional $R^2$ | 0.88 | | | | |
| Marginal $R^2$ | 0.88 | | | | |
| Random effects: |  |  |  |  |  |
| Sample tree | 0.000 |  |  | 0.000 | 0.035 |

<sup>a</sup> Model estimates (±SE) and 95% confidence intervals (CI).

**Table S3.**
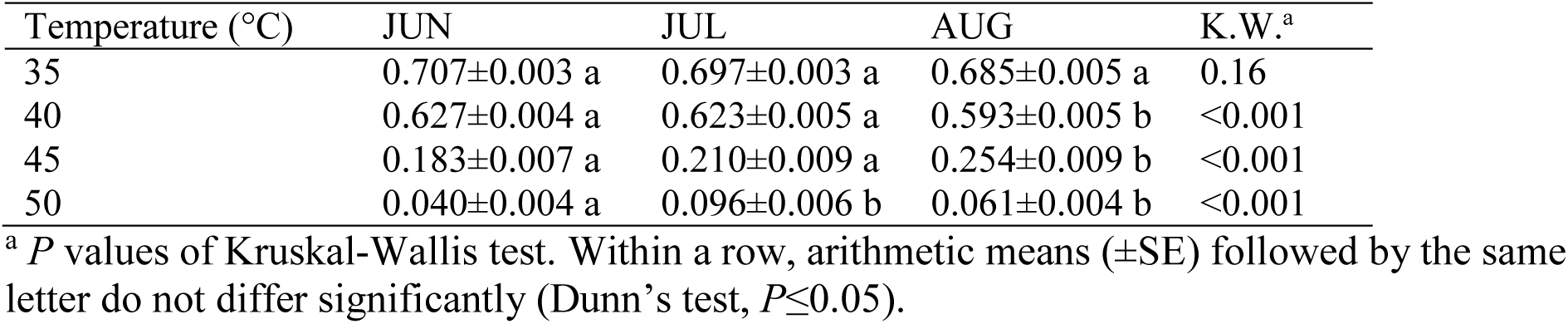
Mean *F_v_/F_m_* in *Larix sibirica* exposed to 35–50°C for 15 min to 4 h in June, July, and August.

**Table S4.**
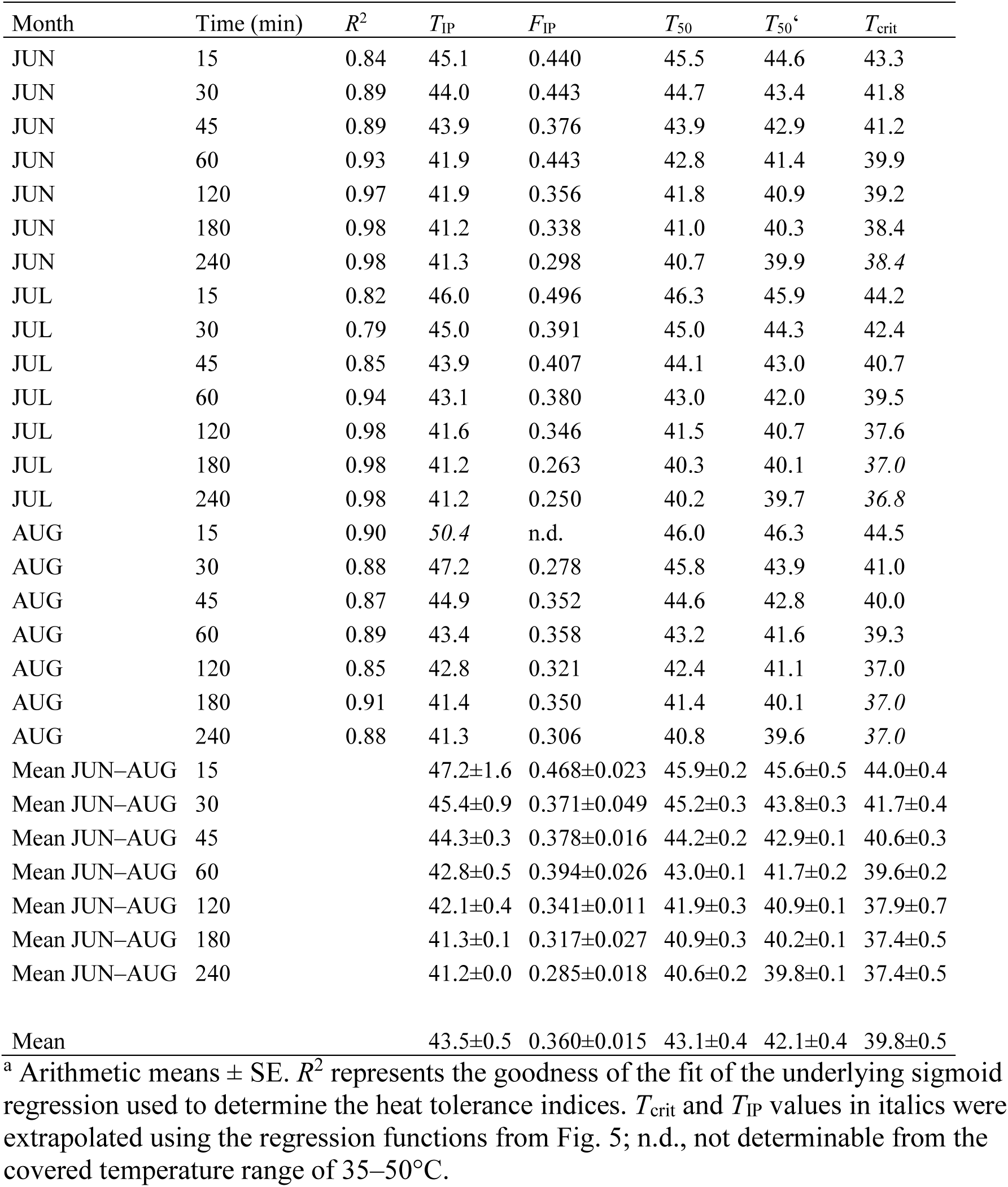
Heat tolerance indices derived from heat exposure experiments with *Larix sibirica* in individual months June to August and incubation times (15 min–4 h)^a^.

**Table S5.** Generalized linear model (GLM) analyzing effects of the month of sampling and incubation time on heat tolerance indices derived from heat exposure experiments with *Larix sibirica* from June to August 2025^a^.

| Parameter | Estimate | SE | *P* | CI |  |
| --- | --- | --- | --- | --- | --- |
|  |  |  |  | 2.5% | 97.5% |
| *T*<sub>IP</sub>: |  |  |  |  |  |
| (Intercept) | 0.0232 | 0.0001 | <0.001 | 0.0230 | 0.0235 |
| Month | −0.0002 | 0.0001 | 0.05 | −0.0005 | −0.0000 |
| Incubation time | 0.0008 | 0.0001 | <0.001 | 0.0006 | 0.0011 |
| *R*<sup>2</sup> | 0.77 |  |  |  |  |
| *F*<sub>IP</sub>: |  |  |  |  |  |
| (Intercept) | 2.8418 | 0.0778 | <0.001 | 2.6979 | 3.0047 |
| Month | 0.1713 | 0.0715 | 0.03 | 0.0036 | 0.3115 |
| Incubation time | 0.3899 | 0.0868 | <0.001 | 0.2285 | 0.5709 |
| *R*<sup>2</sup> | 0.65 |  |  |  |  |
| *T*<sub>50</sub>: |  |  |  |  |  |
| (Intercept) | 0.0234 | 0.0001 | <0.001 | 0.0231 | 0.0234 |
| Month | 0.0001 | 0.0001 | 0.17 | −0.0003 | −0.0001 |
| Incubation time | 0.0011 | 0.0001 | <0.001 | 0.0009 | 0.0012 |
| *R*<sup>2</sup> | 0.89 |  |  |  |  |
| *T*<sub>50</sub>' |  |  |  |  |  |
| (Intercept) | 0.0238 | 0.0001 | <0.001 | 0.0236 | 0.0240 |
| Month | −0.0001 | 0.0001 | 0.53 | −0.0003 | 0.0001 |
| Incubation time | 0.0011 | 0.0001 | <0.001 | 0.0008 | −0.0013 |
| *R*<sup>2</sup> | 0.79 |  |  |  |  |
| *T*<sub>crit</sub>: |  |  |  |  |  |
| (Intercept) | 0.0248 | 0.0002 | <0.001 | 0.0245 | 0.0251 |
| Month | 0.0004 | 0.0002 | 0.04 | 0.0001 | 0.0007 |
| Incubation time | 0.0015 | 0.0002 | <0.001 | 0.0011 | 0.0018 |
| Month × Incubation time | 0.0006 | 0.0002 | 0.01 | 0.0002 | 0.0009 |
| *R*<sup>2</sup> | 0.86 |  |  |  |  |

<sup>a</sup> Model estimates (±SE) and 95% confidence intervals (CI) based on Gaussian distribution and inverse link function.

**Table S6.** Mean and maximum air temperatures (±SD) at 1.5 m height in the studied *Larix sibirica* stands from June to August 2025.

|  | JUN | JUL | AUG<sup>a</sup> | JUN–AUG<sup>b</sup> |
| --- | --- | --- | --- | --- |
| Mean temperature | 14.9±0.3 | 15.4±2.8 | 17.0±2.0 | 15.4±2.9 |
| Mean maximum temperature | 22.5±4.0 | 22.5±3.5 | 27.2±4.3 | 23.2±4.1 |
| Absolute temperature maximum | 28.4±4.0 | 29.0±3.5 | 30.8±4.3 | 30.8±4.1 |

<sup>a</sup> Only means of 1 to 11 August (heat tolerance measurements completed by then); June and July means of the whole months.
<sup>b</sup> Mean values of daily means of the whole sampling interval (1 June to 11 August 2025).

**Fig. S1.**
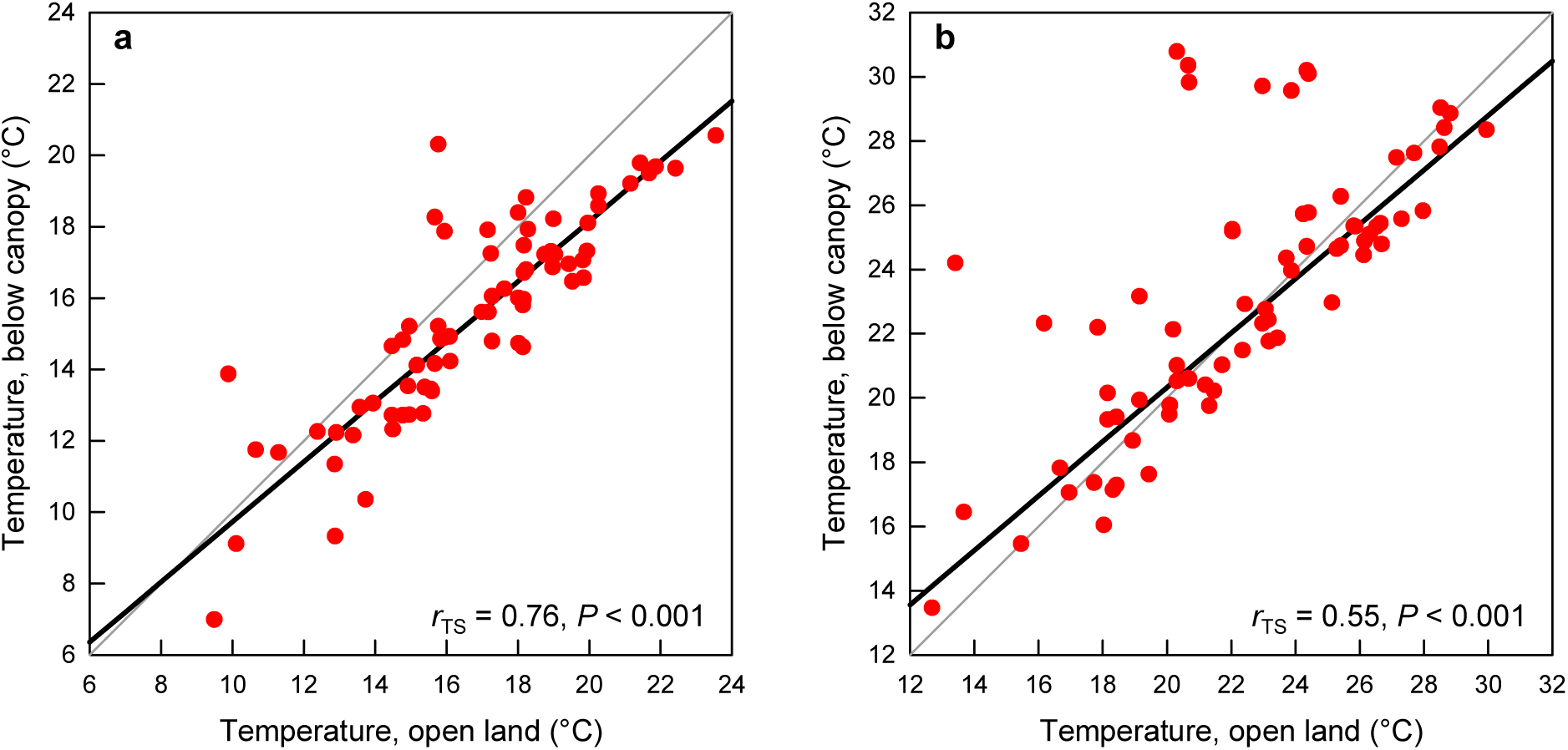
Theil-Sen robust regressions of temperature time series from 1 JUN – 11 AUG 2025 representing (a) daily mean and (b) daily maximum temperatures of hourly data. Open land data are ERA5 reanalysis data for the grid field of 48.5 – 49.0 °N and 100.5 – 101.0 °E modeled for 2 m height. Below-canopy data are measured values recorded at 1.5 m height on our sample plots of *Larix sibirica* forests (*r*_TS_: correlation coefficients from Theil-Sen regression).

**Fig. S2.**
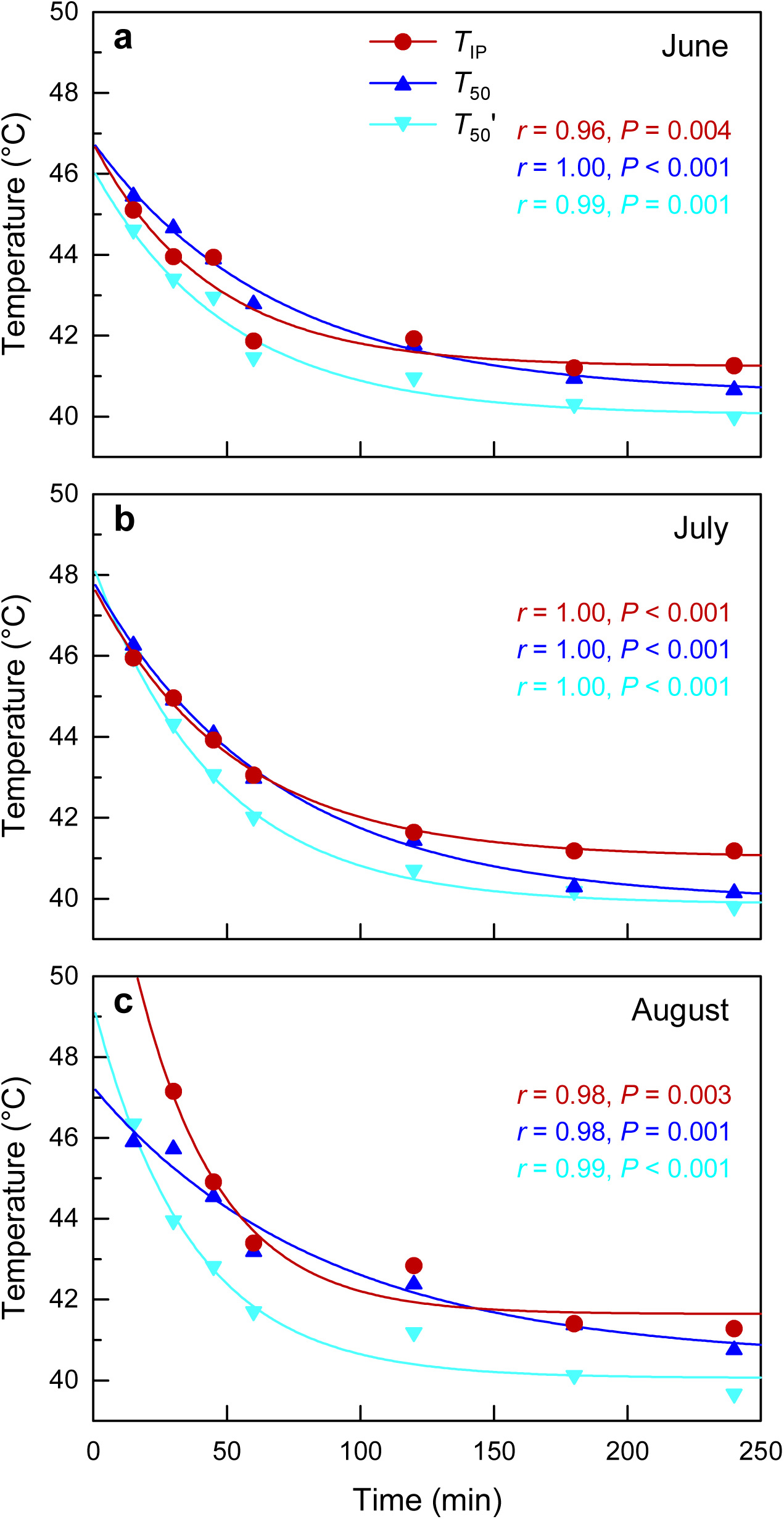
Comparison of different heat tolerance indices *T*_IP_, *T*_50_, and *T*_50_’ vs. incubation time in *Larix sibirica* determined in (a) June, (b) July, and (c) August 2025.

**Fig. S3.**
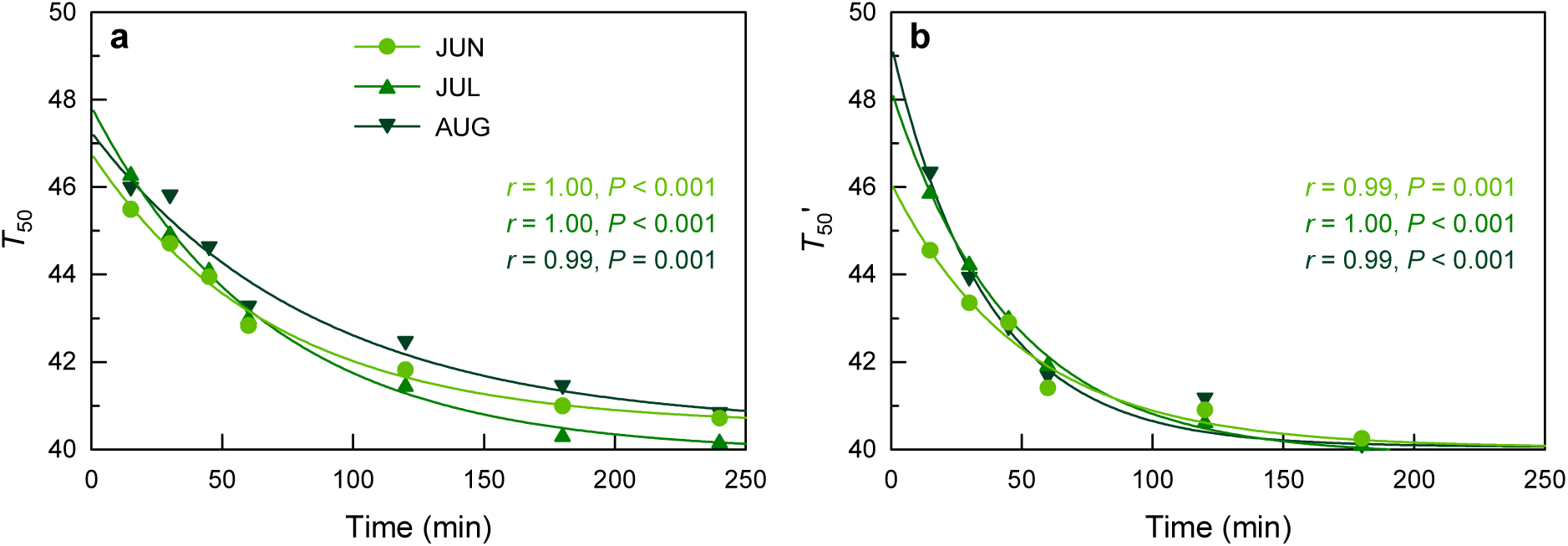
Comparison of different heat tolerance indices: Change of (a) *T*_50_ and (b) *T*_50_’ vs. incubation time in *Larix sibirica* from June to August 2025.

**Fig. S4.**
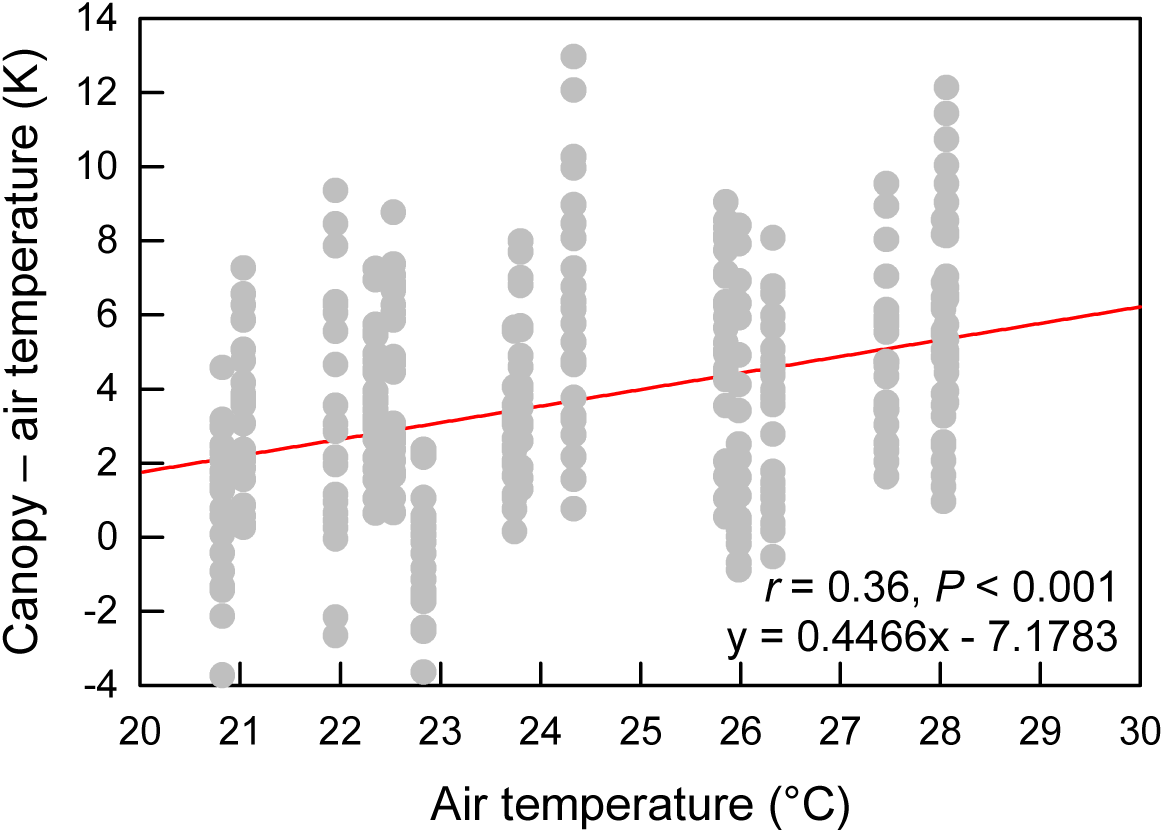
Difference of canopy minus air temperature in relation to air temperature at the time of measurement. Individual drone flights for thermal imaging of canopy temperature share the same air temperature.

## References

Allakhverdiev SI, Kreslavski VD, Klimov VV, Los DA, Carpentier R, Mohanty P (2008) Heat stress: an overview of molecular responses in photosynthesis. Photosynth Res 98:541–550

Allen CD, Macalady AK, Chenchouni H et al. (2010) A global overview of drought and heat-induced tree mortality reveals emerging climate change risks for forests. For Ecol Manag 259:660–684

Anderegg WRL, Hicke JA, Fisher RA et al. (2015) Tree mortality from drought, insects, and their interactions in a changing climate. New Phytol 208:674–683

Anderson-Teixeira KJ, Herrmann V, Rollinson CR (2022) Joint effects of climate, tree size, and year on annual tree growth derived from tree-ring records of ten globally distributed forests. Glob Change Biol 28:245–266

Berry J, Björkman O (1980) Photosynthetic response and adaptation to temperature in higher plants. Ann Rev Plant Physiol 31:491–543

Breshears DD, Adams HD, Eamus D, McDowell NG, Law DJ, Will RE, Williams AP, Zou CB (2013) The critical amplifying role of increasing atmospheric moisture demand on tree mortality and associated regional die-off. Front Plant Sci 4:266

Cailleret M, Dakos V, Jansen S et al. (2019) Early-warning signals of individual tree mortality based on annual radial growth. Front Plant Sci 9:1964.

Caldeira MC (2019) The timing of drought coupled with pathogens may boost tree mortality. Tree Physiol 39:1–5

Cansler CA, Wright MC, vanMantgem PJ, Shearman TM, Varner JM, Hood SM (2024) Drought before fire increases tree mortality after fire. Ecosphere15:e70083

Curtis EM, Knight CA, Petrou K, Leigh A (2014) A comparative analysis of photosynthetic recovery from thermal stress: a desert plant case study. Oecologia 175:1051–1061

Dagvadorj D, Natsagdorj L, Dorjpurev J, Namkhainyam B (2009) Mongolia: assessment report on climate change 2009. Ministry of Nature, Enviroment and Tourism, Ulan Bator, Mongolia

Da Silva BHP, Rossatto DR (2024) Leaf tolerance to heat is independent of leaf phenology in neotropical savanna trees. Trees 38:1343–1350

Didion-Gency M, Gauthey A, Johnson KM, Schuler P, Grossiord C (2025) Leaf excision and exposure duration alter the estimates of the irreversible photosynthetic thermal thresholds. Plant Cell Environ 48:5357–5368

Dulamsuren C (2026) Elastic adjustment in a drought-tolerant boreal conifer compared to a broadleaved pioneer. Agric For Meteorol 388:111323

Dulamsuren C, Hauck M (2008) Spatial and seasonal variation of climate on steppe slopes of the northern Mongolian mountain taiga. Grassland Sci 54:217–230

Dulamsuren C, Hauck M (2021) Drought stress mitigation by nitrogen in boreal forests inferred from stable isotopes. Glob Change Biol 27:5211–5224

Dulamsuren C, Hauck M, Bader M, Osokhjargal D, Oyungerel Sh, Nyambayar S, Runge M, Leuschner C (2009) Water relations and photosynthetic performance in *Larix sibirica* growing in the forest-steppe ecotone of northern Mongolia. Tree Physiol 29:99–110

Dulamsuren C, Hauck M, Leuschner C (2010) Recent drought stress leads to growth reductions in *Larix sibirica* in the western Khentey, Mongolia. Glob Change Biol 16:3024–3035

Dulamsuren C, Bat-Enerel B, Meyer P, Leuschner C (2022) Did stand opening 60 years ago predispose a European beech population to death? Trees For People 8:100265

Dulamsuren C, Byamba-Yondon G, Oyungerel S, Nitschke R, Gebauer T (2023a) Non-structural carbohydrate concentrations in contrasting dry and wet years in early- and late-successional boreal forest trees. Trees 37:1315–1332

Endris J, Rehm E (2025) Leaf temperatures exceed thermal heat tolerances for a community of eastern North America hardwood trees, AoB Plants 17:plae060

Garab G, Yaguzhinsky LS, Dlouhý O, Nesterov SV, Špunda V, Gasanoff ES (2022) Structural and functional roles of non-bilayer lipid phases of chloroplast thylakoid membranes and mitochondrial inner membranes. Progr Lipid Res 86:101163

Gauthey A, Kahmen A, Limousin J-M, Vilagrosa A, Didion-Gency M, Mas E, Milano A, Tunas A, Grossiord C (2024) High heat tolerance, evaporative cooling, and stomatal decoupling regulate canopy temperature and their safety margins in three European oak species. Glob Change Biol 30:e17439

Gounaris K, Brain ARR, Quinn PJ, Williams WP (1983) Structural and functional changes associated with heat-induced phase separations of non-bilayer lipids in chloroplast thylakoid membranes. FEBS Lett 153:47–52

Gounaris K, Brain ARR, Quinn PJ, Williams WP (1984) Structural reorganisation of chloroplast thylakoid membranes in response to heat-stress. Biochim Biophys Acta 766:198–208

Gu L, Grodzinski B, Han J, Marie T, Zhang Y-J, Song YC, Sun Y (2022) Granal thylakoid structure and function: explaining an enduring mystery of higher plants. New Phytol 236:319–329

Haldimann P, Feller U (2005) Growth at moderately elevated temperature alters the physiological response of the photosynthetic apparatus to heat stress in pea (*Pisum sativum* L.) leaves. Plant Cell Environ 28:302–317

Hammond WM, Williams AP, Abatzoglou JT, Adams HD, Klein T, López R, Sáenz-Romero C, Hartmann H, Breshears DD, Allen CD (2022) Global field observations of tree die-off reveal hotter-drought fingerprint for Earth’s forests. Nat Commun 13:1761

Harris I, Osborn TJ, Jones P, Lister D (2020) Version 4 of the CRU TS monthly high-resolution gridded multivariate climate dataset. Sci Data 7:109

Hauck M, Dulamsuren C (2026) T_50_ as a proxy of PS II heat tolerance: time dependence, differences in calculation methods and implications for its suitability to assess the heat tolerance of tree species. bioRxiv 2026.07.24.740464. 10.64898/2026.07.24.740464

Hauck M, Dulamsuren C, Leuschner C (2016) Anomalous increase in winter temperature and decline in forest growth associated with severe winter smog in the Ulan Bator basin. Water Air Soil Pollut 227:261

Hauck M, Schneider T, Bahlinger S, Fischbach J, Oswald G, Csapek G, Dulamsuren C (2025) Heat tolerance of temperate tree species from Central Europe. For Ecol Manag 580:122541

Hauck M, Csapek G, Krämer K, Schmidt O, Lucas Y, Popp L, Linda Szafranek L, Dulamsuren C (2026) Heat tolerance and its seasonal acclimation in *Fagus sylvatica* compared to *Fagus orientalis* and *Pseudotsuga menziesii*. bioRxiv 2026.05.17.725742. 10.64898/2026.05.17.725742

Havaux M (1993) Characterization of thermal damage to the photosynthetic electron transport system in potato leaves. Plant Sci 94:19–33

Húdoková H, Petrik P, Petek-Petrik A, Konôpková A, Leštianka A, Střelcová K, Kmet J, Kurjak D (2022) Heat-stress response of photosystem II in five ecologically important tree species of European temperate forests. Biologia 77:671–680

Hüve K, Bichele I, Rasulov B, Niinemets Ü (2011) When it is too hot for photosynthesis: heat-induced instability of photosynthesis in relation to respiratory burst, cell permeability changes and H_2_O_2_ formation. Plant Cell Environ 34:113–126

Ji S, Yang S, Al-Sakkaf AS, Seka AM, Zhang J (2025) Intensified compound drought and heat events in typical arid-semiarid transition region over the last four decades based on a daily-scale index. Sci Total Environ 1001:180541

Keen RM, Voelker SL, Wang S-YS, Bentz BJ, Goulden ML, Dangerfield CR, Reed CC, Hood SM, Csank AZ, Dawson TE, Merschel AG, Still CJ (2022) Changes in tree drought sensitivity provided early warning signals to the California drought and forest mortality event. Glob Change Biol 28:1119–1132

Khansaritoreh E, Dulamsuren C, Klinge M, Ariunbaatar T, Bat-Enerel B, Batsaikhan G, Ganbaatar K, Saindovdon D, Yeruult Y, Tsogtbaatar J, Tuya D, Leuschner C, Hauck M (2017) Higher climate warming sensitivity of Siberian larch in small than large forest islands in the fragmented Mongolian forest steppe. Glob Change Biol 23:3675–3689

Klinge M, Dulamsuren C, Erasmi S, Hauck M (2018) Climate effects on vegetation vitality at the treeline of boreal forests of Mongolia. Biogeosciences 15:1319–1333

Krause GH, Winter K, Krause B, Jahns P, García M, Aranda J, Virgo A (2010) High-temperature tolerance of a tropical tree, *Ficus insipida*: methodological reassessment and climate change considerations. Funct Plant Biol 37:890–900

Kreslavski V, Tatarinzev N, Shabnova N, Semenova G, Kosobryukhov A (2008) Characterization of the nature of photosynthetic recovery of wheat seedlings from short-term dark heat exposures and analysis of the mode of acclimation to different light intensities. J Plant Physiol 165:1592–1600.10.1016/j.jplph.2007.12.011

Kunert N, Hajek P (2022) Shade-tolerant temperate broad-leaved trees are more sensitive to thermal stress than light-demanding species during a moderate heatwave. Trees For People 9:100282

Kunert N, Hajek P, Hietz P, Morris H, Rosner S, Tholen D (2022) Summer temperatures reach the thermal tolerance threshold of photosynthetic decline in temperate conifers. Plant Biol 24:1254–1261

Kunert N, Ehrmann J, Gebhard S, Hofmann S, Zimmermann G, Hajek P (2026a) Temperate tree species show cross-tolerance to heat, drought, and late spring-frost stress. New Phytol. 10.1111/nph.71277

Kunert N, Düsterhöft E, Stumpf B (2026b) Heat treatment duration affects in vitro-induced photosynthetic impairment and development of necrotic leaf tissue in three Mediterranean oak species. Plant Biol 28:1256–1267

Leuzinger S, Körner C (2007) Tree species diversity affects canopy leaf temperatures in a mature temperate forest. Agric For Meteorol 146:29–27

Lípová L, Krchňák P, Komenda J, Ilík P (2010) Heat-induced disassembly and degradation of chlorophyll-containing protein complexes in vivo. Biochim Biophys Acta Bioenerg 1797:63–70

Liu H, Williams AP, Allen CD, Guo D, Wu X, Anenkhonov OA, Liang EY, Sandanov DV, Yin Y, Qi Z, Badmaeva NK (2013) Rapid warming accelerates tree growth decline in semi-arid forests of Inner Asia. Glob Change Biol 19:2500–2510

Liu H, Shangguan H, Zhou M, Airebule P, Zhao P, He W, Xiang C, Wu X (2019) Differentiated responses of nonstructural carbohydrate allocation to climatic dryness and drought events in the Inner Asian arid timberline. Agric For Meteorol 271:355–361

Mamet SD, Chun KP, Metsaranta JM, Barr AG, Johnstone JF (2015) Tree rings provide early warning signals of jack pine mortality across a moisture gradient in the southern boreal forest. Environ Res Lett 10:084021

Mantova M, Menezes-Silva PE, Badel E, Cochard H, Torres-Ruiz JM (2021) The interplay of hydraulic failure and cell vitality explains tree capacity to recover from drought. Physiol Plant 172:247–257

Mantova M, Cochard H, Burlett R, Delzon S, King A, Rodriguez-Dominguez CM, Ahmed MA, Trueba S, Torres-Ruiz JM (2023) On the path from xylem hydraulic failure to downstream cell death. New Phytol 237:793–806

Mirabel A, Girardin MP, Metsaranta J, Way D, Reich PB (2023) Increasing atmospheric dryness reduces boreal forest tree growth. Nat Commun 14:6901

Mohanty P, Kreslavski VD, Klimov VV, Los DA, Mimuro M, Carpentier R, Allakhverdiev SI (2012) Heat stress: susceptibility, recovery and regulation. In: Eaton-Rye JJ, Tripathy BC, Sharkey TD (eds) Photosynthesis. Plastid biology, energy conversion and carbon assimilation. Springer, Dordrecht, pp 251–274

Münchinger I, Hajek P, Akdogan B, Caicoya AT, Kunert N (2023) Leaf thermal tolerance and sensitivity of temperate tree species are correlated with leaf physiological and functional drought resistance traits. J For Res 34:63–76

Murata N, Takahashi S, Nishiyama Y, Allakhverdiev SI (2007) Photoinhibition of photosystem II under environmental stress. Biochim Biophys Acta Bioenerget 1767:414–421

Murchie EH, Lawson T (2013) Chlorophyll fluorescence analysis: a guide to good practice and understanding some new applications. J Exp Bot 64:3983–3998

Neuner G, Buchner O (2023) The dose makes the poison: The longer the heat lasts, the lower the temperature for functional impairment and damage. Environ Exp Bot 212:105395

O’Sullivan OS, Heskel MA, Reich PD et al. (2017) Thermal limits of leaf metabolism across biomes. Glob Change Biol 23:209–223

Perez TM, Feeley KJ (2020) Photosynthetic heat tolerances and extreme leaf temperatures. Funct Ecol 34:2236–2245

Perez TM, Socha A, Tserej O, Feeley KJ (2021). Photosystem II heat tolerances characterize thermal generalists and the upper limit of carbon assimilation. Plant Cell Environ 44:2321–2330

Petrik P, Petek-Petrik A, Konôpková A, Fleischer P, Stojnic S, Zavadilova I, Kurjak D (2023) Seasonality of PSII thermostability and water use efficiency of in situ mountainous Norway spruce (*Picea abies*). J For Res 34:197–208

Posch BC, Amoanimaa-Dede, H, Aparecido LMT, Atkin OK et al. (2025) High-temperature acclimation of photosystem II in land plants. New Phytol 249:1108–1123

Pospíšil P (2016) Production of reactive oxygen species by photosystem II as a response to light and temperature Stress. Front Plant Sci 7:1950

Rennenberg H, Loreto F, Polle A, Brilli F, Fares S, Beniwal RS, Gessler A (2006) Physiological responses of forest trees to heat and drought. Plant Biol 8: 556–571

Reszczyńska E, Hanaka A (2020) Lipids composition in plant membranes. Cell Biochem Biophys 78:401–414

Richter R, Hutengs C, Wirth C, Bannehr L, Vohland M (2021) Detecting tree species effects on forest canopy temperatures with thermal remote sensing: the role of spatial resolution. Remote Sens 13:135

Salvucci ME, Crafts-Brandner SJ (2004) Inhibition of photosynthesis by heat stress: the activation state of Rubisco as a limiting factor in photosynthesis. Physiol Plant 120:179–186

Slot M, Cala D, Aranda J, Virgo A, Michaletz ST, Winter K (2021) Leaf heat tolerance of 147 tropical forest species varies with elevation and leaf functional traits, but not with phylogeny. Plant Cell Environ 44:2414–2427

Still, CJ, Powell R, Aubrecht D, Kim Y, Helliker B, Roberts D, Richardson AD, Goulden M (2019) Thermal imaging in plant and ecosystem ecology: applications and challenges. Ecosphere 10:e02768

Still CJ, Rastogi B, Page G, Griffith DM, Sibley A, Schulze M, Hawkins L, Pau S, Detto M, Helliker BR (2021) Imaging canopy temperature: shedding (thermal) light on ecosystem processes. New Phytol 230:1746–1753

Still CJ, Page G, Rastogi B et al. (2022) No evidence of canopy-scale leaf thermoregulation to cool leaves below air temperature across a range of forest ecosystems. Proc Natl Acad Sci USA 119:e2205682119

Subbotina A, Chernyak E, Soukhovolsky V, Morozov S, Martemyanov V (2025) Latitudinal variation in constitutive chemical defense compounds in two host plants of *Lymantria dispar* (Lymantriidae): *Betula pendula* (Betulaceae) and *Larix sibirica* (Pinaceae). For Ecol Manag 590:122811

Tiwari R, Gloor E, da Cruz WJA et al. (2021) Photosynthetic quantum efficiency in south-eastern Amazonian trees may be already affected by climate change. Plant Cell Environ 44:2428–2439

Tripathy KP, Mukherjee S, Mishra AK, Mann ME, Williams AP (2023) Climate change will accelerate the high-end risk of compound drought and heatwave events. Proc Natl Acad Sci USA 120:e2219825120

van Tiel N, Lenczner G, Rao MP, Grossiord C, Tuia D (2026) Tropical forests are facing increasing risks of exposure to critical temperature thresholds. Proc Natl Acad Sci USA 123:e2528622123

Williams AP, Allen CD, Macalady AK et al. (2013) Temperature as a potent driver of regional forest drought stress and tree mortality. Nat Clim Change 3:292–297

Yamamoto Y (2016) Quality control of photosystem II: the mechanisms for avoidance and tolerance of light and heat stresses are closely linked to membrane fluidity of the thylakoids. Front Plant Sci 7:1136

Yamamoto Y, Aminaka R, Yoshioka M, Khatoon M, Komayama K, Takenaka D, Yamashita A, Nijo N, Inagawa K, Morita N, Sasaki T, Yamamoto Y (2008) Quality control of photosystem II: impact of light and heat stresses. Photosynth Res 98:589–608

Yamashita A, Nijo N, Pospíšil P, Morita N, Takenaka D, Aminaka R, Yamamoto Y, Yamamoto Y (2008) Quality control of photosystem II: reactive oxygen species are responsible for the damage to photosystem II under moderate heat stress. J Biol Chem 283:28380–28391

Yi K, Smith JW, Jablonski AD, Tatham EA, Scanlon TM, Lerdau MT, Novick KA, Yang X (2020) High heterogeneity in canopy temperature among co-occurring tree species in a temperate forest. J Geophys Res Biogeosci 125:e2020JG005892

Zheng G, Tian B, Zhang F, Tao F, Li W (2011) Plant adaptation to frequent alterations between high and low temperatures: remodelling of membrane lipids and maintenance of unsaturation levels. Plant Cell Environ 34:1431–1442

Zhu L, Scafaro AP, Vierling E, Ball MC, Posch BC, Stock F, Atkin OK (2024) Heat tolerance of a tropical–subtropical rainforest tree species *Polyscias elegans*: time-dependent dynamic responses of physiological thermostability and biochemistry. New Phytol 241:715–731

